# Effects of different proportions of ground corn and steam-flaked corn diets on growth performance, serum biochemical parameters, rumen fermentation and rumen microbial metagenome of yaks

**DOI:** 10.1101/2025.09.08.675018

**Authors:** Zhang Xi-rui, Hu Zhi-bin, Peng Zhong-li, Wang Hai-bo, Huang Yan-ling

## Abstract

This study objected to assess the effect of substituting ground corn (GC) with steam-flaked corn (SFC) at various ratios on growth performance, serum biochemistry, rumen fermentation, and microbiome in fattening yaks. Fifty male Maiwa yaks (196.43 ± 16.08 kg) were randomly assigned to five groups: SFC 0, SFC 25, SFC 50, SFC 75, and SFC 100, representing 0%, 25%, 50%, 75%, and 100% substitution of GC with SFC. Results showed that the dry-matter intake (DMI) in SFC 0 group was significantly lower than those in other treatment groups (*P* < 0.05). Among the serum biochemical parameters, the triglyceride (TG) content in the SFC 100 group was significantly higher than other treatment groups (*P* < 0.05), with a quadratically increase (*P* < 0.05). Regarding rumen fermentation parameters, the propionate in the SFC 50 and SFC 100 groups was significantly higher than in the other groups (*P* < 0.05). At the phylum level of the rumen microbiome, the replacing with SFC significantly increased the relative abundance of *Planctomycetota*, *Bacteria*, and *Candidatus_Saccharibacteria* (*P* < 0.05). At the genus level, the replacing with SFC significantly increased the relative abundance of *Paludibacteraceae* and *Bacteria* (*P* < 0.05). Among the CAZy enzymes in the rumen, the relative abundance of cellulase modules (CM) in the SFC 0 group was significantly higher than other groups (*P* < 0.05). In conclusion, replacing 50% of GC with SFC may be an effective strategy to enhances growth performance in yak farming by improving rumen fermentation, microbiome composition, and carbohydrate-active enzymes.

**Highlights:**

- The replacement of GC with SFC can optimize rumen fermentation of yaks.
- The replacement of GC with SFC in the diets affected the rumen microbiome of yaks.
- Metagenome profiled differences between CAZy enzymes and eggNOG functional proteins.

## 1. Introduction

The Qinghai-Tibet Plateau, as the largest and highest plateau on Earth, is located at the center of the Asian continent, covering an area of approximately 2.5 million square kilometers with an average elevation of around 5,000 meters (Fang et al., 2024). It is characterized by extreme climatic conditions, including cold temperatures, low oxygen levels, and intense ultraviolet radiation (Chen et al., 2022). This region is considered one of the most inhospitable areas for habitation on earth. Its vast expanse and extreme environment present significant challenges for the survival of both plants and animals. However, the yak (*Bos grunniens*) has successfully adapted to this harsh environment, becoming one of the few large animals capable of surviving year-round in this region(Ayalew et al., 2021;Ding et al., 2020;Malekkhahi et al., 2021;Xin et al., 2019). Yaks are ancient mammals with physiological characteristics distinct from other ruminants(Ahmad et al., 2024). Their exceptional adaptability to extreme high-altitude environments, including their cold tolerance and ability to thrive in low-oxygen conditions, makes them ideal companions for the highland Tibetan people, providing them with invaluable resources for survival (Huang et al., 2021). Yak farming is one of the key industries in the pastoral areas of western Sichuan, offering a primary source of income for local residents and playing a crucial role in the stability and development of the regional economy (Gao et al., 2022;Long et al., 1999;Wang et al., 2011). In recent years, with the rapid development of animal husbandry in western Sichuan, significant breakthroughs have been made in yak farming techniques. However, challenges have also emerged. The question of how to adopt sustainable farming practices that protect the fragile ecological environment of the plateau, while promoting the yak industry and preventing overgrazing and grassland degradation, is critical to ensuring the long-term development of the industry. This requires the rational utilization of feed resources in large-scale intensive farming, ensuring production performance while reducing feed costs to improve the efficiency of yak farming.

Energy is a crucial component of ruminant diets, and the adequacy of energy supply directly influences animal growth, development, and production performance (F. N. Owens. et al., 1993) Selecting appropriate energy sources and effectively managing energy supply can help optimize diet formulation, reduce protein requirements, and minimize nitrogen emissions by providing sufficient energy. Utilizing suitable feed processing techniques, such as grinding, pelleting, and steam pelleting, can enhance nitrogen utilization in feed. By designing diets that achieve nitrogen balance, it is possible to meet the nutritional needs of ruminants while also reducing negative environmental impacts, thereby contributing to the sustainable development of livestock farming (Itzhak Mizrahi et al., 2021).

Corn plays an important role in animal feed ingredients globally, due to its nutritional value, availability, and economic importance. Rich in starch, corn provides a major energy source for both poultry and livestock In livestock production, blending corn processed through various methods can maximize its nutritional value while ensuring that animals receive a balanced and comprehensive diet (SALIM HM et al., 2010). Different processing methods can affect the nutritional properties of corn to varying degrees, improving its digestibility, energy utilization, and protein availability. Mixing corn processed by different techniques can compensate for the potential nutritional deficiencies of a single processing method, achieving a more balanced feed formulation(CORONA et al., 2005;Lee et al., 2023). Furthermore, this approach allows for flexible adjustments in feed formulations based on practical needs, resulting in cost savings and enhanced production efficiency and economic benefits in livestock farming(Buttrey et al., 2013). In this experiment, steam-flaked corn was used to replace ground corn in the fattening yak diet at five levels: 0%, 25%, 50%, 75%, and 100%. The study aimed to investigate the effects of these substitutions on yak growth performance, serum biochemical parameters, rumen fermentation characteristics, and rumen microbial metagenomics. The findings will provide insights into the optimal inclusion ratio of steam-flaked corn in fattening yak diets, offering a scientific reference for the application of steam-flaked corn in yak husbandry.

## 2. Materials and methods

### 2.1. Animals, diets and experimental design

All procedures in this study were approved by the Animal Care and Use Committee of Southwest Minzu University (Protocol No. SMU 202106010). The experiment was conducted at Kangbala Green Food Co., Ltd. in Ganzi County, Ganzi Prefecture, Sichuan Province (100°02′E, 31°61′N, approximately 3000 meters above sea level). During the experimental period, the temperature in the housing facility ranged from 5 to 14 °C. The experiment was designed as a completely randomized design with a single factor. Fifty male Maiwa yaks, approximately 4 years old, with similar body conditions (196.43 ± 16.08 kg), were selected as the experimental animals. The trial was divided into 5 groups, with 10 replicates per group, each consisting of one yak. The five treatment groups (SFC 0, SFC 25, SFC 50, SFC 75, and SFC 100) were fed with steam-flaked corn replacing ground corn in the basal diet at proportions of 0%, 25%, 50%, 75%, and 100%, respectively. Prior to the experiment, all pens were disinfected and cleaned. Each yak received deworming treatment and vaccination against common infectious diseases. After a 25-day adaptation period, the experiment lasted for 75 days. The experimental diet was primarily composed of concentrate and silage, with a concentrate-to-forage ratio of 55:45. The composition of the diet ingredients and their nutritional components are presented in Table 1. A total mixed ration (TMR) was mixed and fed fresh. The yaks were fed twice daily at 8:00 and 16:00, with free access to both feed and water.

**Table. 1.**
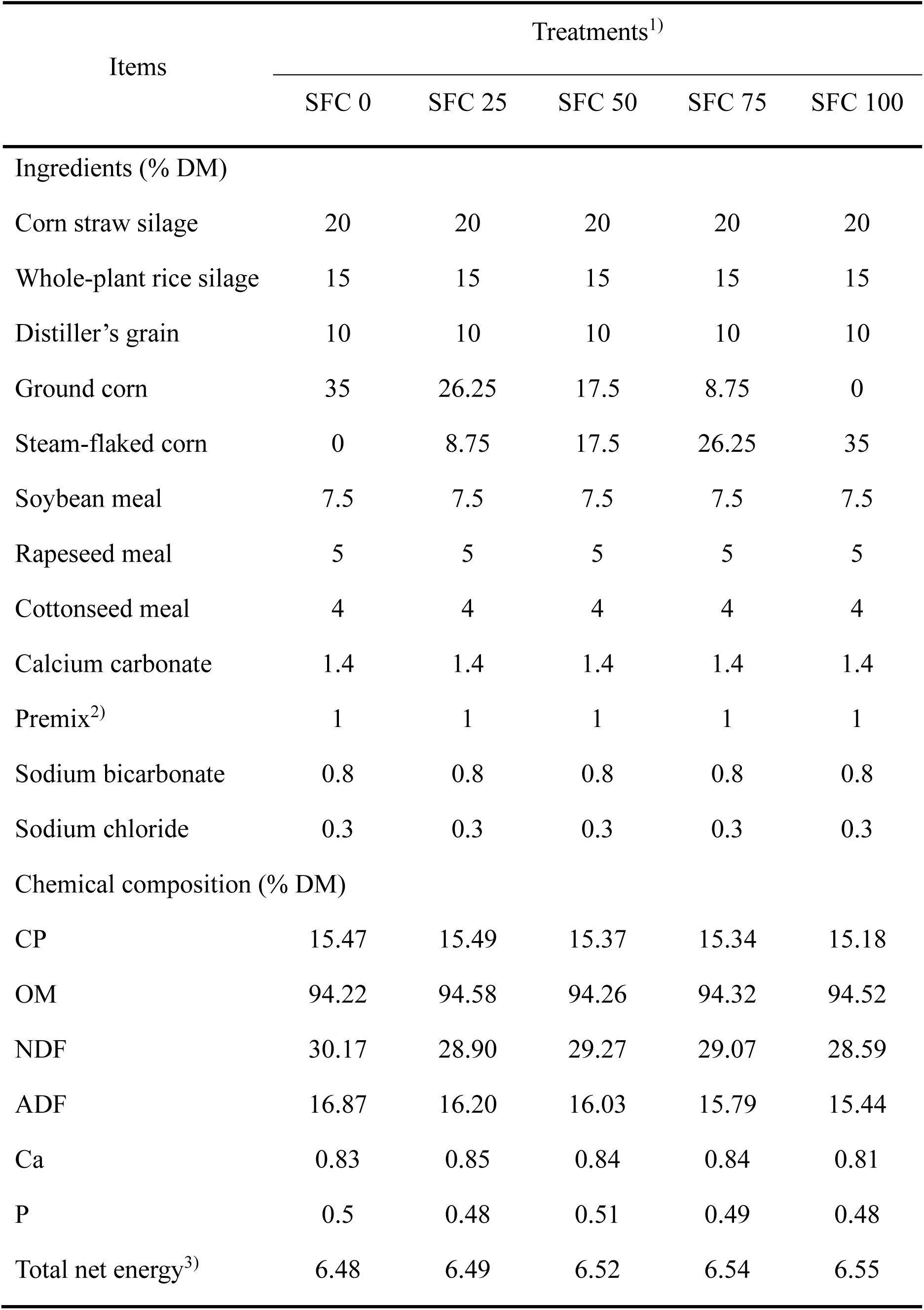
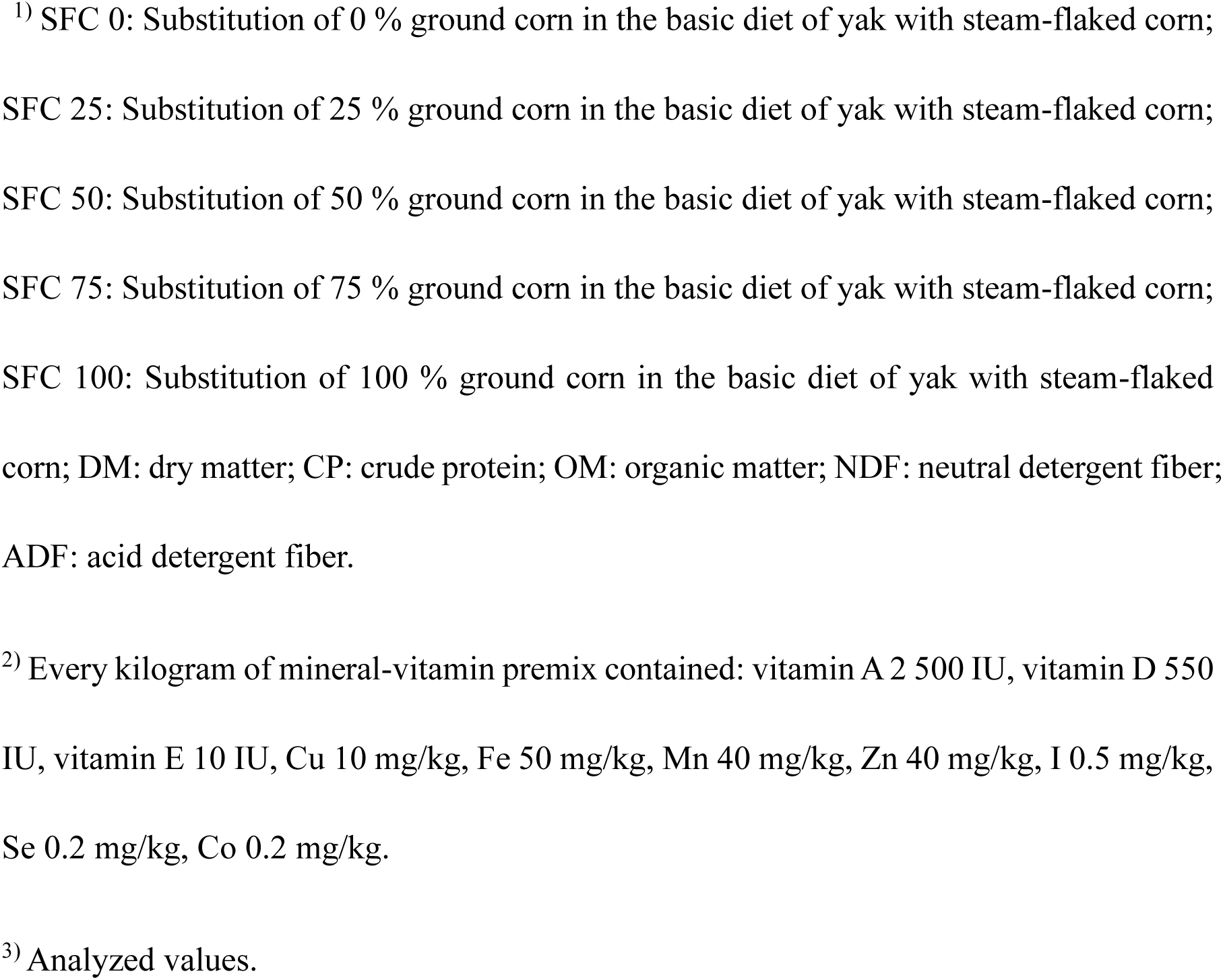
Experimental diet ingredients and nutrition composition (DM, %).

| Items | Treatments<sup>1)</sup> |  |  |  |  |
| --- | --- | --- | --- | --- | --- |
|  | SFC 0 | SFC 25 | SFC 50 | SFC 75 | SFC 100 |
| Ingredients (% DM) |  |  |  |  |  |
| Corn straw silage | 20 | 20 | 20 | 20 | 20 |
| Whole-plant rice silage | 15 | 15 | 15 | 15 | 15 |
| Distiller's grain | 10 | 10 | 10 | 10 | 10 |
| Ground corn | 35 | 26.25 | 17.5 | 8.75 | 0 |
| Steam-flaked corn | 0 | 8.75 | 17.5 | 26.25 | 35 |
| Soybean meal | 7.5 | 7.5 | 7.5 | 7.5 | 7.5 |
| Rapeseed meal | 5 | 5 | 5 | 5 | 5 |
| Cottonseed meal | 4 | 4 | 4 | 4 | 4 |
| Calcium carbonate | 1.4 | 1.4 | 1.4 | 1.4 | 1.4 |
| Premix<sup>2)</sup> | 1 | 1 | 1 | 1 | 1 |
| Sodium bicarbonate | 0.8 | 0.8 | 0.8 | 0.8 | 0.8 |
| Sodium chloride | 0.3 | 0.3 | 0.3 | 0.3 | 0.3 |
| Chemical composition (% DM) |  |  |  |  |  |
| CP | 15.47 | 15.49 | 15.37 | 15.34 | 15.18 |
| OM | 94.22 | 94.58 | 94.26 | 94.32 | 94.52 |
| NDF | 30.17 | 28.90 | 29.27 | 29.07 | 28.59 |
| ADF | 16.87 | 16.20 | 16.03 | 15.79 | 15.44 |
| Ca | 0.83 | 0.85 | 0.84 | 0.84 | 0.81 |
| P | 0.5 | 0.48 | 0.51 | 0.49 | 0.48 |
| Total net energy<sup>3)</sup> | 6.48 | 6.49 | 6.52 | 6.54 | 6.55 |

<sup>1)</sup> SFC 0: Substitution of 0 % ground corn in the basic diet of yak with steam-flaked corn; SFC 25: Substitution of 25 % ground corn in the basic diet of yak with steam-flaked corn; SFC 50: Substitution of 50 % ground corn in the basic diet of yak with steam-flaked corn; SFC 75: Substitution of 75 % ground corn in the basic diet of yak with steam-flaked corn; SFC 100: Substitution of 100 % ground corn in the basic diet of yak with steam-flaked corn; DM: dry matter; CP: crude protein; OM: organic matter; NDF: neutral detergent fiber; ADF: acid detergent fiber.
<sup>2)</sup> Every kilogram of mineral-vitamin premix contained: vitamin A 2 500 IU, vitamin D 550 IU, vitamin E 10 IU, Cu 10 mg/kg, Fe 50 mg/kg, Mn 40 mg/kg, Zn 40 mg/kg, I 0.5 mg/kg, Se 0.2 mg/kg, Co 0.2 mg/kg.
<sup>3)</sup> Analyzed values.

### 2.2. Sample collection

On Day 1 and Day 76 of the experiment, the yaks were weighed after a 12-hour fast to record their initial body weight (IBW) and final body weight (FBW). The average daily gain (ADG) was then calculated. During the experiment, feed intake and leftovers were recorded at each feeding, and the average daily feed intake (ADFI) was calculated based on the amount of TMR provided and the remaining feed. Feed conversion ratio (FCR) was calculated as the ratio of ADG to ADFI.

On the morning of Day 76, prior to feeding, six yaks were randomly selected from each group. A 20 mL blood sample was drawn from the jugular vein of each yak and placed in a vacuum blood collection tube. After collection, the samples were immediately centrifuged at 3494×g for 10 minutes at 4°C. The supernatant was then aliquoted into 2 mL cryovials and stored in liquid nitrogen for further analysis.

Rumen fluid samples were collected using an oral catheter method, with a disposable syringe used to aspirate the fluid. Between each animal, the catheter was thoroughly cleaned with water. To minimize potential contamination from saliva, the first 50 mL of rumen fluid was discarded, and approximately 60 mL of rumen fluid was then collected and aliquoted into cryovials. The samples were stored in liquid nitrogen to preserve their biological activity for subsequent analysis.

### 2.3. Nutrient Analysis

Feed samples were ground to pass through a 1 mm sieve. The dry matter (DM), crude protein (CP), calcium (Ca), phosphorus (P), and ash content of the feed were determined according to the procedures outlined by AOAC (Hasan, 2015). Acid detergent fiber (ADF) and neutral detergent fiber (NDF) were measured using the Van Soest method with filter bags (Van Soest et al., 1991).

### 2.4. Serum Biochemical Parameters

After thawing the samples at 4°C, serum levels of alanine aminotransferase (ALT), aspartate aminotransferase (AST), albumin (ALB), lactate dehydrogenase (LDH), alkaline phosphatase (ALP), triglycerides (TG), total cholesterol (TC), total protein (TP), glucose (GLU), creatine kinase (CK), creatinine (CREA), urea (UREA), and pyruvate (PYR) were determined using Hitachi 3100 (Hitachi Co., Tokyo, Japan). The concentration of globulin (GLB) was calculated by subtracting ALB from TP.

### 2.5. Rumen Fermentation Parameter

The concentration of NH_3_-N was determined using the alkaline phenol hypochlorite method (Weatherburn, 1967). Microbial crude protein (MCP) was quantified using a bicinchoninic acid kit (Solarbio Life Sciences, Beijing, China) following the procedure outlined in a previous study (Zhang et al., 2022). Volatile fatty acids (VFA) were analyzed by gas chromatography (Agilent 6890N, Santa Clara, CA, USA) equipped with a DB-WAXETR column (30 m × 1.0 μm × 0.53 mm, Agilent, USA), as described in our previous method (Zhang, et al. 2022). In brief, 2-ethylbutyric acid (2EB) served as an internal standard, and quantification was performed using the internal standard correction method. The operational parameters were as follows: the carrier gas was nitrogen, the split ratio was 40:1, the flow rate was 2.0 mL/min, the average linear velocity was 38 cm/s, and the column pressure was 11.3 psi. The temperature program started at 120 °C (held for 3 minutes) and was increased to 180 °C (held for 1 minute) at a rate of 10 °C/min. The flame ionization detector (FID) temperature was set at 250 °C. Hydrogen flow was 40 mL/min, air flow was 45 mL/min, column flow was 45 mL/min, the inlet temperature was 210 °C, and the injection volume was 0.6 μL.

### 2.6. Metagenomic sequencing and analysis

Genomic DNA was extracted from rumen liquid samples using the FastDNA® Spin Kit for Soil (MP Biomedicals, California, U.S.) following the manufacturer’s instructions. DNA concentration and purity were measured with TBS-380 and NanoDrop2000, respectively, and DNA quality was assessed on a 1% agarose gel. DNA was fragmented to about 400 bp using Covaris M220 (Gene Company Limited, China) for paired-end library construction, employing NEXTFLEX Rapid DNA-Seq (Bioo Scientific, Austin, TX, USA). Adapters with sequencing primer sites were ligated to fragment ends. Paired-end sequencing was performed on Illumina NovaSeq (Illumina Inc., San Diego, CA, USA) at Majorbio Bio-Pharm Technology Co., Ltd. (Shanghai, China) using NovaSeq Reagent Kits (www.illumina.com).The raw reads have been submitted to the National Center for Biotechnology Information (NCBI) Sequence Read Archive (SRA) database under Accession Number: PRJNA953596 (available at http://www.ncbi.nlm.nih.gov/sra, accessed on 5 April 2023).

Data analysis was conducted on the Majorbio Cloud Platform (www.majorbio.com). Paired-end reads were trimmed for adaptors, and low-quality reads (length < 50 bp, quality < 20, or containing N bases) were removed using fastp (Chen et al., 2019) (https://github.com/OpenGene/fastp, version 0.20.0). Metagenomic data assembly was done using MEGAHIT (Li et al. 2015) (https://github.com/voutcn/megahit, version 1.1.2), selecting contigs ≥ 300 bp for further gene prediction and annotation. Open reading frames (ORFs) were predicted using MetaGene (Noguchi et al., 2006) (http://metagene.cb.k.u-tokyo.ac.jp/), with ORFs ≥ 100 bp translated into amino acid sequences using the NCBI translation table. A non-redundant gene catalog was built using CD-HIT (Fu et al., 2012) (http://www.bioinformatics.org/cd-hit/, version 4.6.1) with 90% sequence identity and coverage. High-quality reads were aligned to the catalog to calculate gene abundance (95% identity) using SOAPaligner (Li et al., 2008) (http://soap.genomics.org.cn/, version 2.21). Representative sequences from the non-redundant gene catalog were aligned to the NR database with an e-value cutoff of 1e^-5^ using Diamond (Buchfink et al., 2015) (http://www.diamondsearch.org/index.php, version 0.8.35) for taxonomic annotations. COG and KEGG annotations were performed using Diamond (Buchfink et al., 2015) (http://www.diamondsearch.org/index.php, version 0.8.35) against the eggNOG (1e^-5^ cutoff) and KEGG (http://www.genome.jp/keeg/, 1e^-5^ cutoff) databases, respectively.

### 2.7. Statistical analysis

Statistical analysis was performed using the general linear model (GLM) procedure of SAS 9.4 (SAS Institute Inc., Cary, NC, USA) for one-way analysis of variance (ANOVA). Multiple comparisons were conducted using the LSD test while orthogonal polynomial analysis was applied to evaluate the linear or quadratic responses of the indicators. Spearman rank correlation analysis was used for correlation analysis. Results are presented as means and pooled SEM. In order to confirm the significant differences, *P* < 0.05 was considered statistically significant and 0.05 ≤ *P* < 0.10 was considered a trend.

## 3. Results

### 3.1. Growth performance

As shown in Table 2, both WG and ADG in each group exhibited a quadratic trend, initially increasing and then decreasing (*P* = 0.094). The DMI of the SFC 0 group was significantly lower than that of the other groups and followed a quadratic decline.

**Table. 2.** Effects of different proportions of GC and SFC diets on growth performance of yaks.

| Items | Treatments<sup>1)</sup> |  |  |  |  | SEM | *P* value<sup>2)</sup> |  |  |
| --- | --- | --- | --- | --- | --- | --- | --- | --- | --- |
|  | SFC 0 | SFC 25 | SFC 50 | SFC 75 | SFC 100 |  | Treatment | Linear | Quadratic |
| IBW (kg) | 203.55 | 209.20 | 203.60 | 206.10 | 200.40 | 18.454 | 0.877 | 0.618 | 0.513 |
| FBW (kg) | 283.65 | 295.75 | 298.35 | 299.20 | 285.90 | 25.415 | 0.528 | 0.753 | 0.083 |
| ADG (kg/d) | 1.07 | 1.15 | 1.26 | 1.24 | 1.14 | 0.255 | 0.432 | 0.360 | 0.094 |
| DMI (kg/d) | 6.96<sup>a</sup> | 7.14<sup>b</sup> | 7.02<sup>b</sup> | 7.14<sup>b</sup> | 6.82<sup>b</sup> | 0.136 | 0.030 | 0.290 | 0.013 |
| FCR | 6.68 | 6.12 | 5.77 | 5.93 | 6.39 | 1.292 | 0.547 | 0.546 | 0.098 |

<sup>1)</sup> SFC 0: Substitution of 0 % ground corn in the basic diet of yak with steam-flaked corn; SFC 25: Substitution of 25 % ground corn in the basic diet of yak with steam-flaked corn; SFC 50: Substitution of 50 % ground corn in the basic diet of yak with steam-flaked corn; SFC 75: Substitution of 75 % ground corn in the basic diet of yak with steam-flaked corn; SFC 100: Substitution of 100 % ground corn in the basic diet of yak with steam-flaked corn; SEM: standard error of mean; IBW: initial body weight; FBW: final body weight; ADG: average daily gain; DMI: dry-matter intake; FCR: feed conversion ratio = DMI/ADG.
<sup>2)</sup> Means with different superscript letters in the same row are significantly different (*p* < 0.05).

### 3.2. Serum biochemical indexes

As shown in Table 3, the serum TG content in the SFC 100 group was significantly higher than in the other groups (*P* < 0.05), exhibiting a quadratic increasing (*P* < 0.05). The serum ALB also demonstrated a quadratic pattern, first increasing and then decreasing (*P* < 0.05). The serum UREA showed a linear increase (*P* < 0.05). The serum GLU in the SFC 50 group tended to be higher than in the other treatment groups (*P* = 0.095), while the serum PYR showed a linear decreasing trend (*P* = 0.061). Compared to the SFC 0 group, the replacement of GC with SFC in the diet had no significant effect on the serum ALT, AST, LDH, ALP, TC, TP, GLO, GLU, CK, or CREA in the yaks (*P* > 0.05).

**Table. 3.**
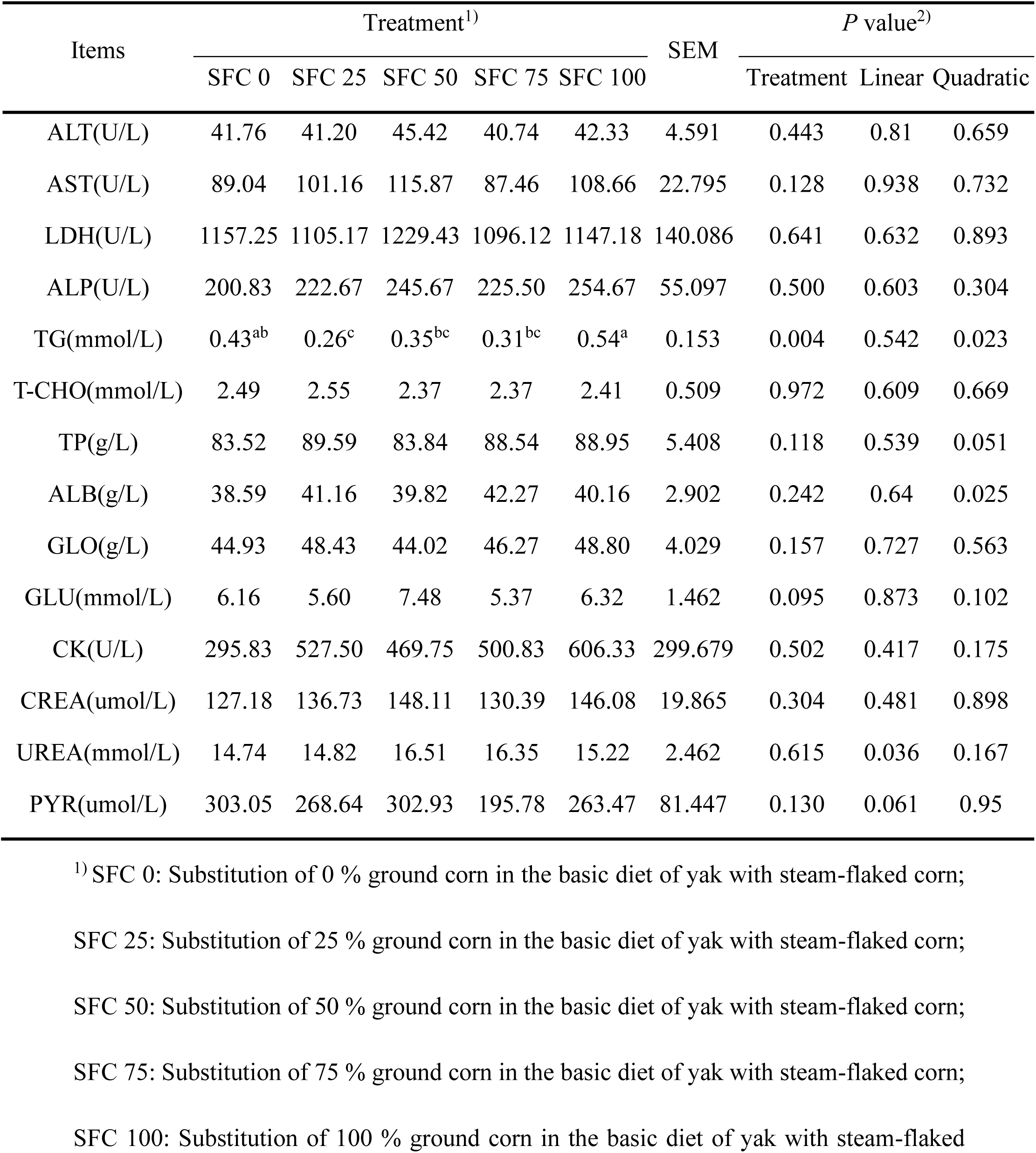
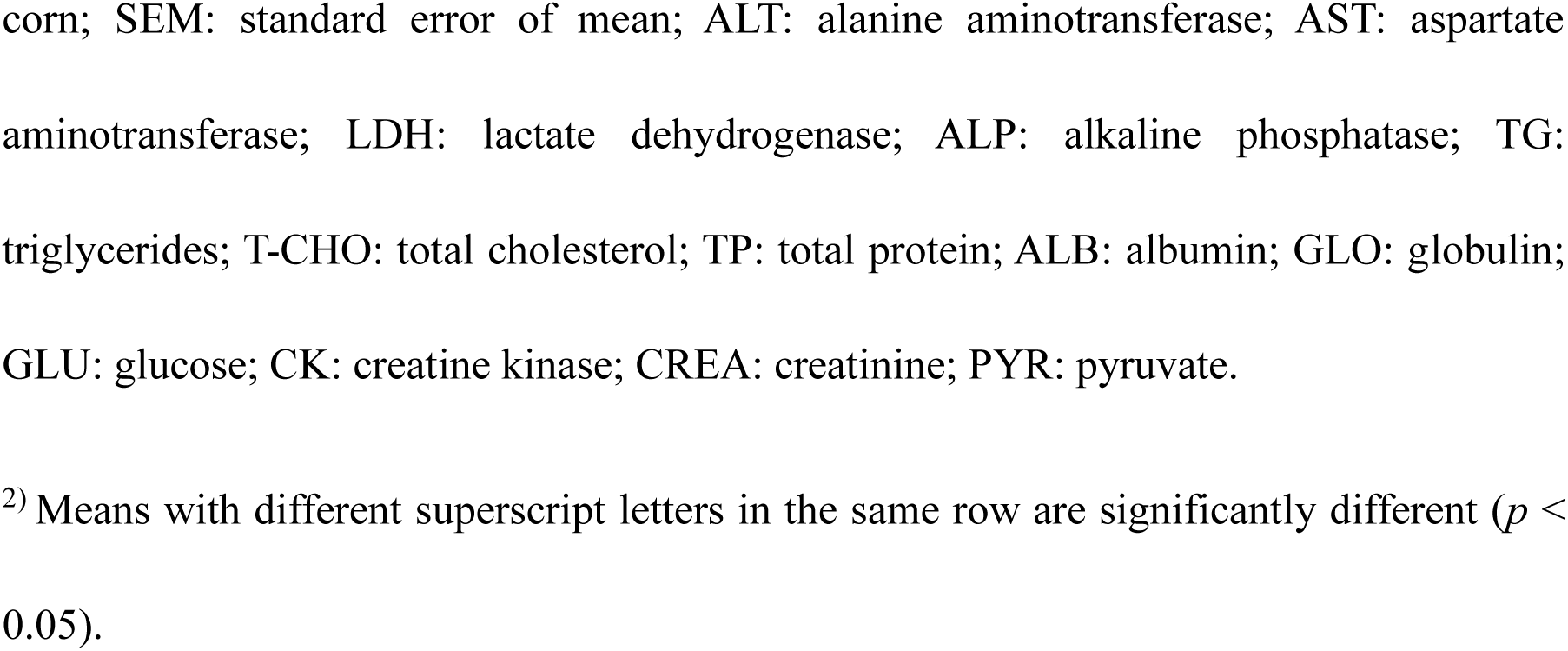
Effects of different proportions of GC and SFC diets on serum biochemical indexes of yaks.

| Items | Treatment<sup>1)</sup> |  |  |  |  | SEM | P value<sup>2)</sup> |  |  |
| --- | --- | --- | --- | --- | --- | --- | --- | --- | --- |
|  | SFC 0 | SFC 25 | SFC 50 | SFC 75 | SFC 100 |  | Treatment | Linear | Quadratic |
| ALT(U/L) | 41.76 | 41.20 | 45.42 | 40.74 | 42.33 | 4.591 | 0.443 | 0.81 | 0.659 |
| AST(U/L) | 89.04 | 101.16 | 115.87 | 87.46 | 108.66 | 22.795 | 0.128 | 0.938 | 0.732 |
| LDH(U/L) | 1157.25 | 1105.17 | 1229.43 | 1096.12 | 1147.18 | 140.086 | 0.641 | 0.632 | 0.893 |
| ALP(U/L) | 200.83 | 222.67 | 245.67 | 225.50 | 254.67 | 55.097 | 0.500 | 0.603 | 0.304 |
| TG(mmol/L) | 0.43<sup>ab</sup> | 0.26<sup>c</sup> | 0.35<sup>bc</sup> | 0.31<sup>bc</sup> | 0.54<sup>a</sup> | 0.153 | 0.004 | 0.542 | 0.023 |
| T-CHO(mmol/L) | 2.49 | 2.55 | 2.37 | 2.37 | 2.41 | 0.509 | 0.972 | 0.609 | 0.669 |
| TP(g/L) | 83.52 | 89.59 | 83.84 | 88.54 | 88.95 | 5.408 | 0.118 | 0.539 | 0.051 |
| ALB(g/L) | 38.59 | 41.16 | 39.82 | 42.27 | 40.16 | 2.902 | 0.242 | 0.64 | 0.025 |
| GLO(g/L) | 44.93 | 48.43 | 44.02 | 46.27 | 48.80 | 4.029 | 0.157 | 0.727 | 0.563 |
| GLU(mmol/L) | 6.16 | 5.60 | 7.48 | 5.37 | 6.32 | 1.462 | 0.095 | 0.873 | 0.102 |
| CK(U/L) | 295.83 | 527.50 | 469.75 | 500.83 | 606.33 | 299.679 | 0.502 | 0.417 | 0.175 |
| CREA(umol/L) | 127.18 | 136.73 | 148.11 | 130.39 | 146.08 | 19.865 | 0.304 | 0.481 | 0.898 |
| UREA(mmol/L) | 14.74 | 14.82 | 16.51 | 16.35 | 15.22 | 2.462 | 0.615 | 0.036 | 0.167 |
| PYR(umol/L) | 303.05 | 268.64 | 302.93 | 195.78 | 263.47 | 81.447 | 0.130 | 0.061 | 0.95 |

<sup>1)</sup> SFC 0: Substitution of 0 % ground corn in the basic diet of yak with steam-flaked corn;
SFC 25: Substitution of 25 % ground corn in the basic diet of yak with steam-flaked corn;
SFC 50: Substitution of 50 % ground corn in the basic diet of yak with steam-flaked corn;
SFC 75: Substitution of 75 % ground corn in the basic diet of yak with steam-flaked corn;
SFC 100: Substitution of 100 % ground corn in the basic diet of yak with steam-flaked
corn; SEM: standard error of mean; ALT: alanine aminotransferase; AST: aspartate aminotransferase; LDH: lactate dehydrogenase; ALP: alkaline phosphatase; TG: triglycerides; T-CHO: total cholesterol; TP: total protein; ALB: albumin; GLO: globulin; GLU: glucose; CK: creatine kinase; CREA: creatinine; PYR: pyruvate.
<sup>2)</sup> Means with different superscript letters in the same row are significantly different ( $p <$ 0.05).

### 3.3. Rumen fermentation parameters

As shown in Table 4, the propionate content in the rumen fluid of the SFC 50 and SFC 100 groups was significantly higher than that of the other groups (*P* < 0.05). With the increasing proportion of SFC replacing GC, the acetic acid and T-VFA contents exhibited a linear increase (*P* < 0.05). No significant differences were observed in the NH_3_-N, MCP, isobutyric acid, butyric acid, isovaleric acid, and valeric acid contents in the rumen fluid of yaks across the experimental groups.

**Table. 4.**
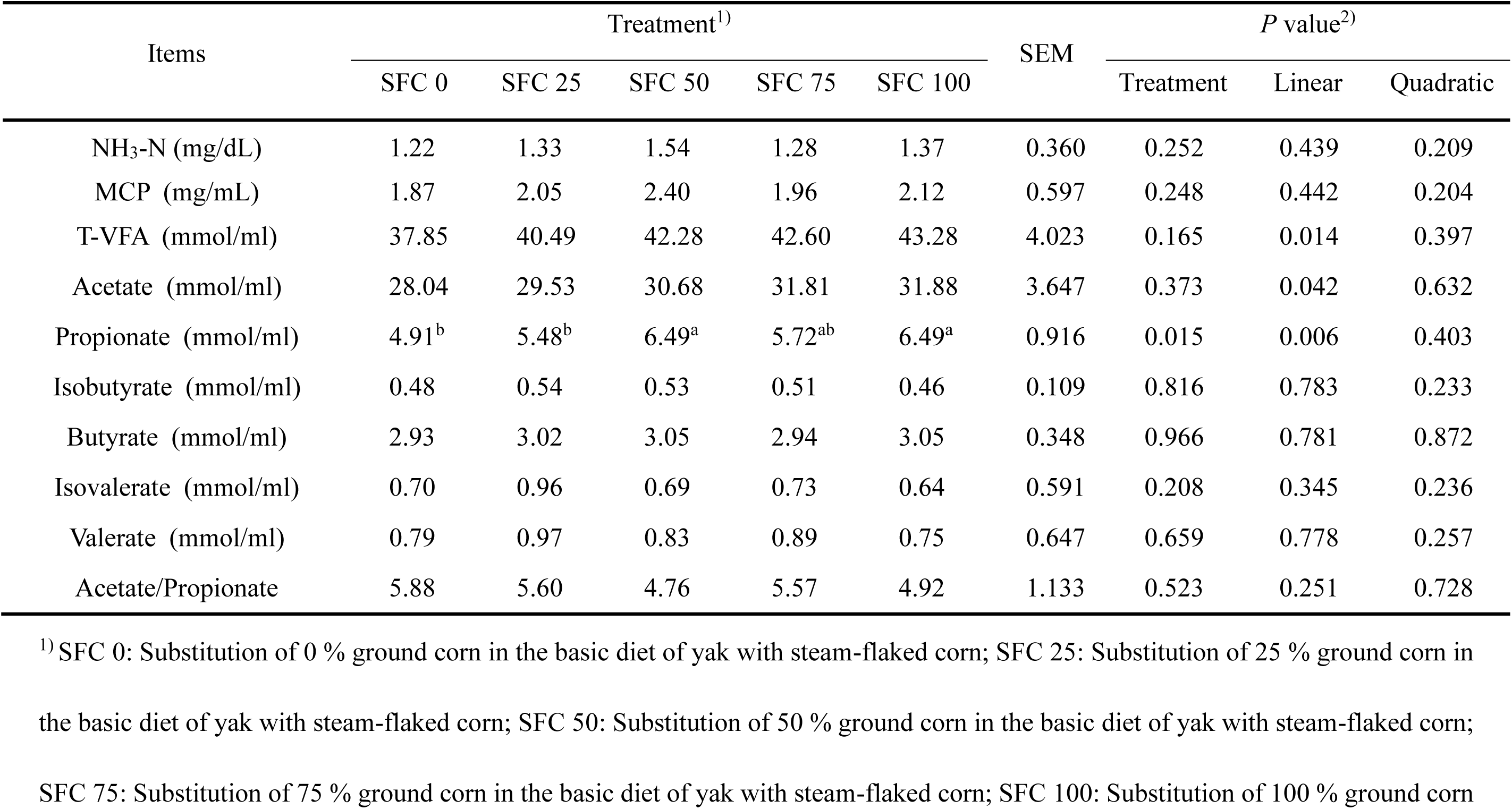
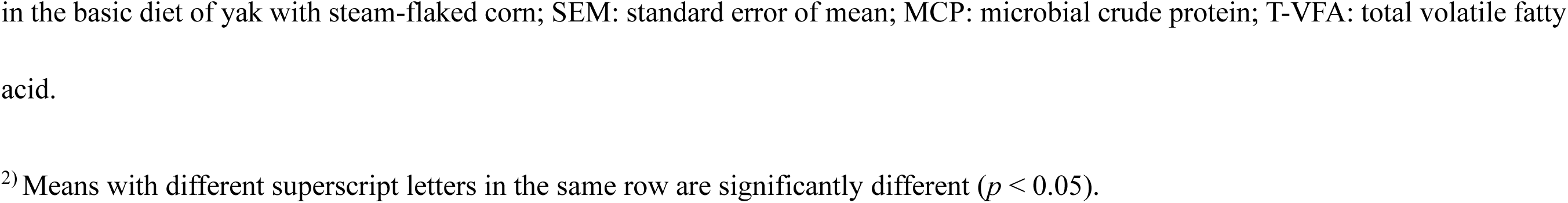
Effects of different proportions of GC and SFC diets on rumen fermentation parameters of yaks.

| Items | Treatment<sup>1)</sup> |  |  |  |  | SEM | *P* value<sup>2)</sup> |  |  |
| --- | --- | --- | --- | --- | --- | --- | --- | --- | --- |
|  | SFC 0 | SFC 25 | SFC 50 | SFC 75 | SFC 100 |  | Treatment | Linear | Quadratic |
| NH<sub>3</sub>-N (mg/dL) | 1.22 | 1.33 | 1.54 | 1.28 | 1.37 | 0.360 | 0.252 | 0.439 | 0.209 |
| MCP (mg/mL) | 1.87 | 2.05 | 2.40 | 1.96 | 2.12 | 0.597 | 0.248 | 0.442 | 0.204 |
| T-VFA (mmol/ml) | 37.85 | 40.49 | 42.28 | 42.60 | 43.28 | 4.023 | 0.165 | 0.014 | 0.397 |
| Acetate (mmol/ml) | 28.04 | 29.53 | 30.68 | 31.81 | 31.88 | 3.647 | 0.373 | 0.042 | 0.632 |
| Propionate (mmol/ml) | 4.91<sup>b</sup> | 5.48<sup>b</sup> | 6.49<sup>a</sup> | 5.72<sup>ab</sup> | 6.49<sup>a</sup> | 0.916 | 0.015 | 0.006 | 0.403 |
| Isobutyrate (mmol/ml) | 0.48 | 0.54 | 0.53 | 0.51 | 0.46 | 0.109 | 0.816 | 0.783 | 0.233 |
| Butyrate (mmol/ml) | 2.93 | 3.02 | 3.05 | 2.94 | 3.05 | 0.348 | 0.966 | 0.781 | 0.872 |
| Isovalerate (mmol/ml) | 0.70 | 0.96 | 0.69 | 0.73 | 0.64 | 0.591 | 0.208 | 0.345 | 0.236 |
| Valerate (mmol/ml) | 0.79 | 0.97 | 0.83 | 0.89 | 0.75 | 0.647 | 0.659 | 0.778 | 0.257 |
| Acetate/Propionate | 5.88 | 5.60 | 4.76 | 5.57 | 4.92 | 1.133 | 0.523 | 0.251 | 0.728 |

<sup>1)</sup> SFC 0: Substitution of 0 % ground corn in the basic diet of yak with steam-flaked corn; SFC 25: Substitution of 25 % ground corn in the basic diet of yak with steam-flaked corn; SFC 50: Substitution of 50 % ground corn in the basic diet of yak with steam-flaked corn; SFC 75: Substitution of 75 % ground corn in the basic diet of yak with steam-flaked corn; SFC 100: Substitution of 100 % ground corn
in the basic diet of yak with steam-flaked corn; SEM: standard error of mean; MCP: microbial crude protein; T-VFA: total volatile fatty acid.
<sup>2)</sup> Means with different superscript letters in the same row are significantly different ( $p < 0.05$ ).

### 3.4. Microbial metagenomic sequence data

Through metagenomic sequencing of rumen samples from yaks, a total of 1,292,975,830 optimized reads were obtained after quality control and host contamination removal, with an average of 46,177,708 reads per sample. Sequence assembly was performed using MEGAHIT v1.1.2 (with the minimum contig length set to ≥300), and the best-assembled sequences were selected for ORF prediction (with gene sequence length ≥100). The assembly yielded a total of 16,639,548 contigs, with an approximate total length of 10.10 Gbp, and 22,539,909 ORFs. After redundancy removal, the number of complete genes was 7,873,724, accounting for 39.71% of the total.

### 3.5. Composition of rumen microbial community

After NR species annotation of the sequencing data, a total of 226 phyla, 462 classes, 987 orders, 2021 families, 5584 genera, and 22,647 species were identified from samples.

Figure 1 shows the top 20 phyla ranked by relative abundance at the phylum level, with *Firmicutes* and *Bacteroidetes* dominating, accounting for 46.97% and 40.88%, respectively. The the abundance of rumen microbial flora in yaks at the phylum level are shown in Table 5. The relative abundance of *Uroviricota* exhibited a quadratic trend, initially increasing and then decreasing (*P* < 0.05). The relative abundance of unclassified*_d Bacteria* in the SFC 75 group was significantly higher than other groups (*P* < 0.05). The relative abundance of *Actinobacteria* showed a quadratic trend, first decreasing and then increasing (*P* < 0.05). The relative abundance of *Planctomycetota* in the SFC 50 group was significantly higher than other groups (*P* < 0.05), exhibiting a quadratic trend of first increasing and then decreasing (*P* < 0.05). The relative abundance of *Candidatus_Saccharibacteria* in the SFC 75 group was significantly higher than other groups (*P* < 0.05), with a quadratic trend of first increasing and then decreasing (*P* < 0.05). The unclassified unclassified*_d Archaea* exhibited a quadratic trend of first increasing and then decreasing (*P* = 0.077).

**Fig. 1.**
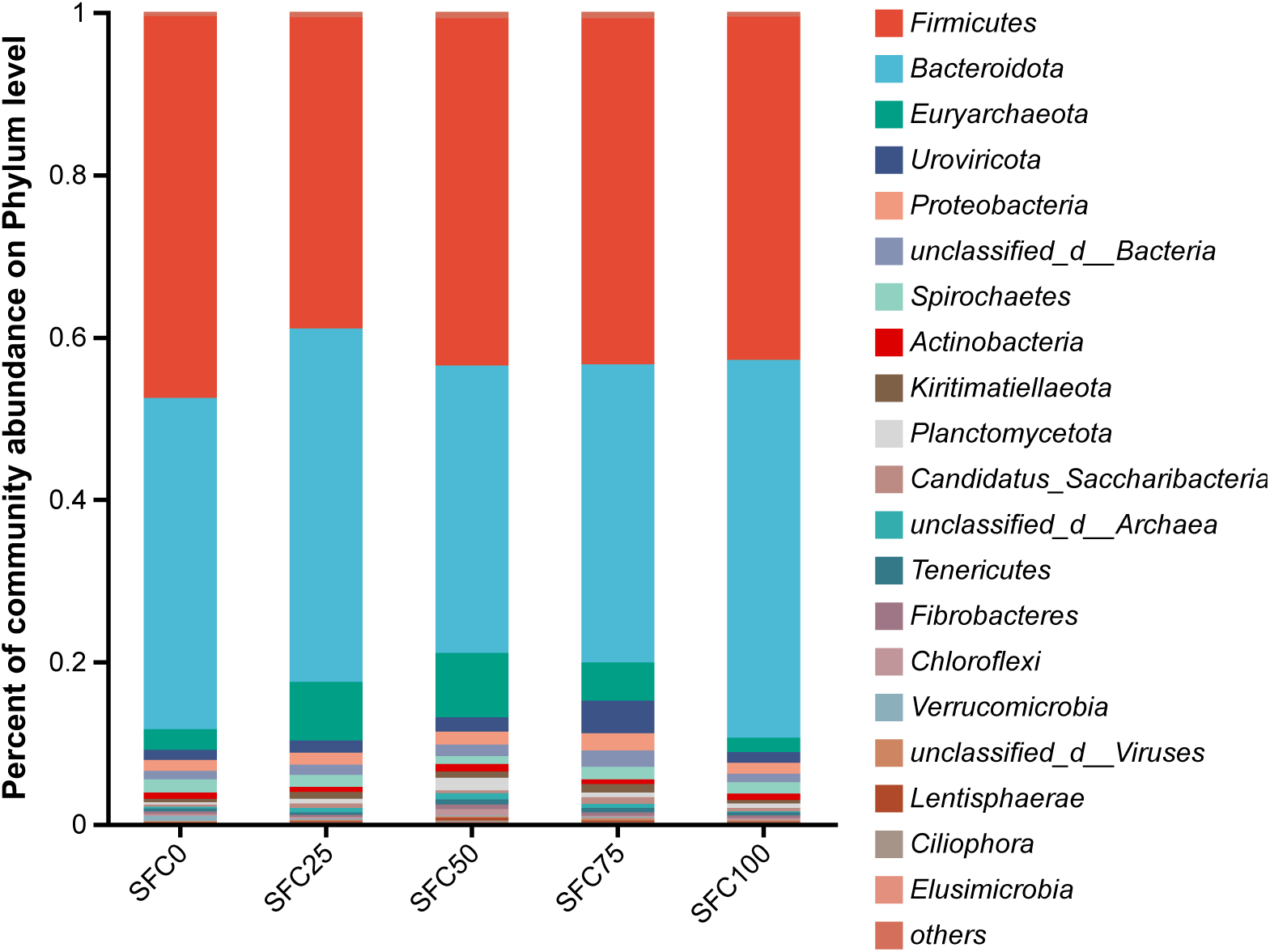
Components of the rumen bacteria in different proportions of GC and SFC diet of yaks. (Phylum level).

**Table. 5.** Effects of different proportions of GC and SFC diets on the abundance of rumen microbial flora in yaks (Phylum, %).

| Items | Treatment<sup>1)</sup> |  |  |  |  | SEM | P value<sup>2)</sup> |  |  |
| --- | --- | --- | --- | --- | --- | --- | --- | --- | --- |
|  | SFC 0 | SFC 25 | SFC 50 | SFC 75 | SFC 100 |  | Treatment | Linear | Quadratic |
| *Firmicutes* | 46.97 | 38.26 | 42.73 | 42.39 | 42.09 | 6.607 | 0.318 | 0.460 | 0.264 |
| *Bacteroidota* | 40.88 | 43.81 | 35.44 | 36.54 | 46.78 | 11.038 | 0.398 | 0.460 | 0.264 |
| *Euryarchaeota* | 2.56 | 7.12 | 7.87 | 4.73 | 1.75 | 4.944 | 0.159 | 0.724 | 0.171 |
| *Uroviricota* | 1.27 | 1.40 | 1.77 | 4.31 | 1.32 | 2.519 | 0.174 | 0.540 | 0.015 |
| *Proteobacteria* | 1.32 | 1.51 | 1.65 | 2.13 | 1.33 | 0.740 | 0.313 | 0.373 | 0.269 |
| unclassified_d_Bacteria | 1.06<sup>b</sup> | 1.27<sup>b</sup> | 1.42<sup>b</sup> | 2.11<sup>a</sup> | 1.12<sup>b</sup> | 0.607 | 0.011 | 0.516 | 0.143 |
| *Spirochaetes* | 1.60 | 1.41 | 0.99 | 1.53 | 1.34 | 0.614 | 0.538 | 0.612 | 0.370 |
| *Actinobacteria* | 0.73 | 0.67 | 0.85 | 0.57 | 0.82 | 0.271 | 0.455 | 0.210 | 0.036 |
| *Kiritimatiellaeota* | 0.45 | 0.84 | 0.83 | 1.07 | 0.41 | 0.563 | 0.200 | 0.849 | 0.623 |
| *Planctomycetota* | 0.29<sup>b</sup> | 0.52<sup>b</sup> | 1.50<sup>a</sup> | 0.59<sup>b</sup> | 0.55<sup>b</sup> | 0.631 | 0.011 | 0.799 | 0.032 |
| *Candidatus_Saccharibacteria* | 0.25<sup>b</sup> | 0.57<sup>ab</sup> | 0.38<sup>b</sup> | 0.81<sup>a</sup> | 0.38<sup>b</sup> | 0.340 | 0.029 | 0.468 | 0.018 |
| unclassified_d_Archaea | 0.23 | 0.55 | 0.80 | 0.49 | 0.20 | 0.521 | 0.302 | 0.223 | 0.077 |

<sup>1)</sup> SFC 0: Substitution of 0 % ground corn in the basic diet of yak with steam-flaked corn; SFC 25: Substitution of 25 % ground corn in the basic diet of yak with steam-flaked corn; SFC 50: Substitution of 50 % ground corn in the basic diet of yak with steam-flaked corn; SFC 75: Substitution of 75 % ground corn in the basic diet of yak with steam-flaked corn; SFC 100: Substitution of 100 % ground corn in the basic diet of yak with steam-flaked corn; SEM: standard error of mean.
<sup>2)</sup> Means with different superscript letters in the same row are significantly different ( $p < 0.05$ ).

Figure 2 displays the top 20 microorganisms ranked by relative abundance at the genus level. Among these, unclassified*_f Bacteroidales*, *Prevotella*, unclassified*_c Clostridia*, unclassified*_f Oscillospiraceae*, and unclassified*_f Lachnospiraceae* dominate, accounting for 20.48%, 12.79%, 12.06%, 10.80%, and 5.12%, respectively. The abundance of the yak rumen microbial flora at the genus level is shown in Table 6. The relative abundance of *Methanobrevibacter* follows a quadratic trend, initially increasing and then decreasing (*P* < 0.05). The relative abundance of unclassified*_f Paludibacteraceae* in the SFC 100 group is significantly higher than that in other treatment groups (*P* < 0.05), with a quadratic trend of first increasing and then decreasing (*P* < 0.05). The relative abundance of unclassified*_d Bacteria* in the SFC 75 group is significantly higher than in other treatment groups, showing a quadratic trend of first increasing and then decreasing (*P* < 0.05). The relative abundance of unclassified*_f Bacteroidales* follows a quadratic trend of first decreasing and then increasing (*P* = 0.073). The relative abundance of unclassified*_c Clostridia* in the SFC 75 group shows a tendency to be significantly higher than in other treatment groups (*P* = 0.063). The relative abundance of unclassified*_f Oscillospiraceae* follows a quadratic trend of first decreasing and then increasing (*P* = 0.061).

**Fig. 2.**
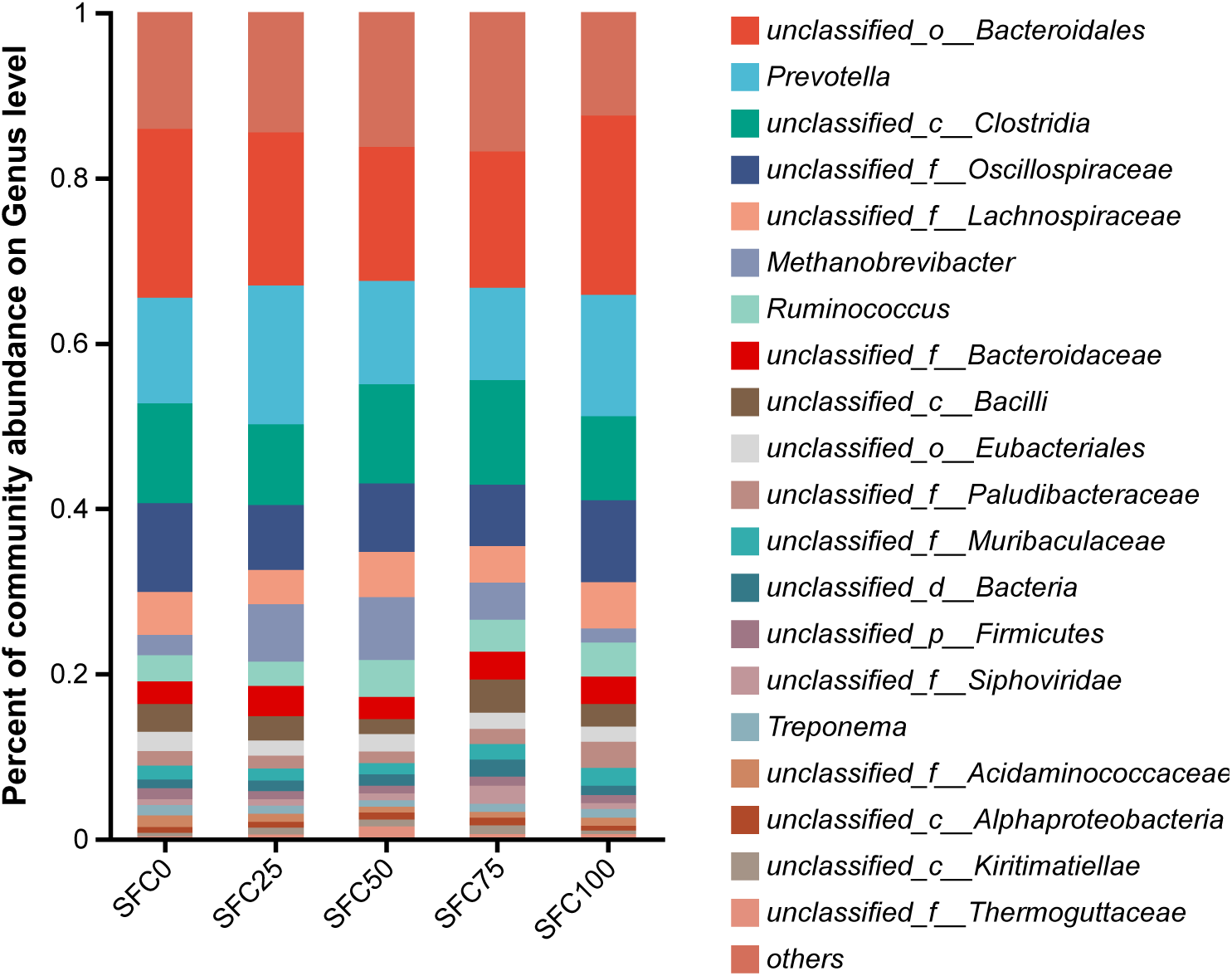
Components of the rumen bacteria in different proportions of GC and SFC diet of yaks. (Genus level).

**Table. 6.**
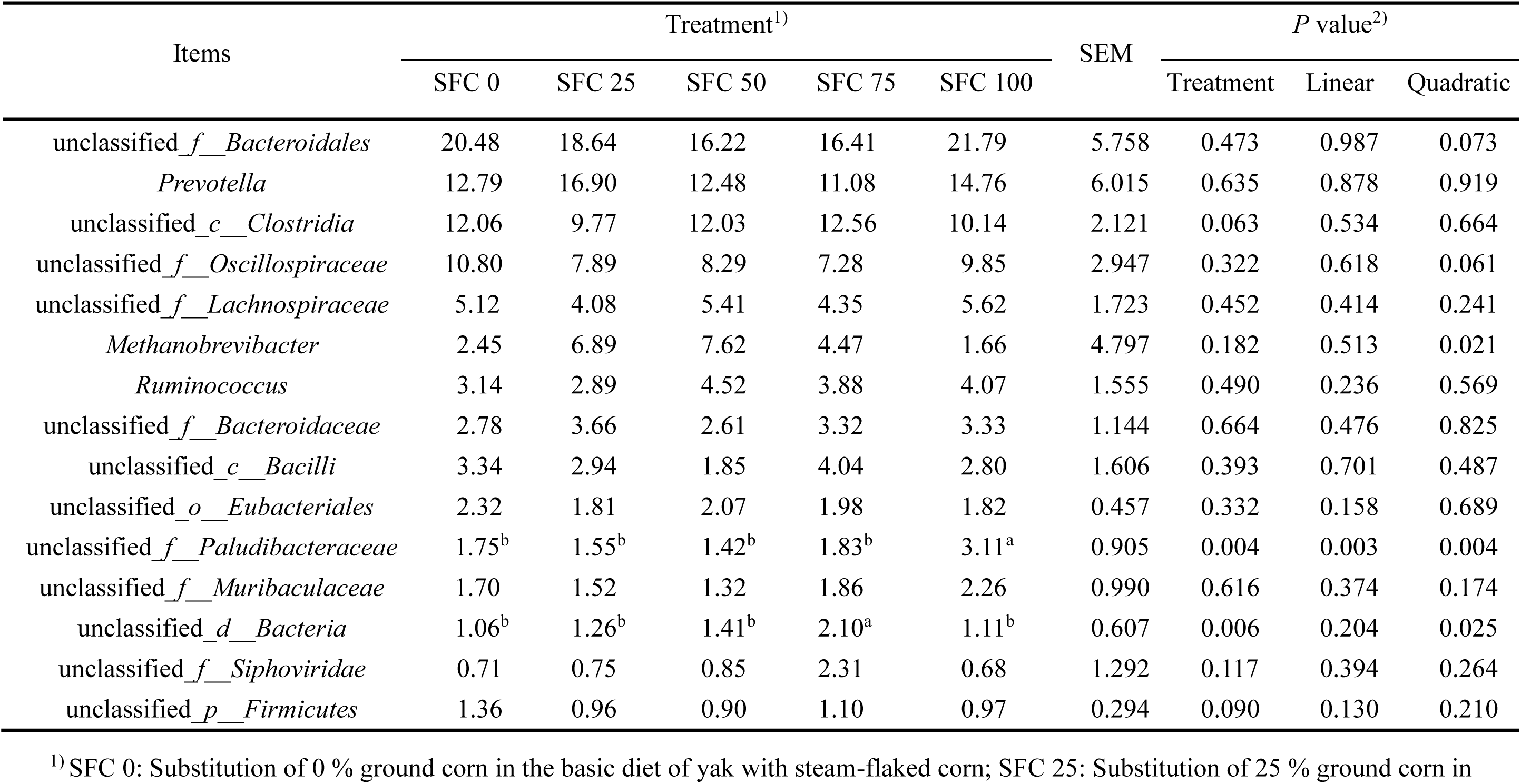
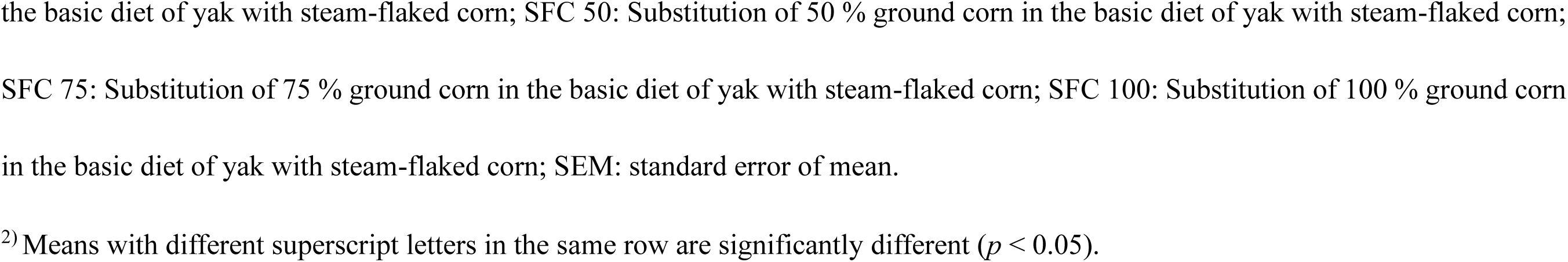
Effects of different proportions of GC and SFC diets on the abundance of rumen microbial flora in yaks (Genus, %).

| Items | Treatment<sup>1)</sup> |  |  |  |  | SEM | P value<sup>2)</sup> |  |  |
| --- | --- | --- | --- | --- | --- | --- | --- | --- | --- |
|  | SFC 0 | SFC 25 | SFC 50 | SFC 75 | SFC 100 |  | Treatment | Linear | Quadratic |
| unclassified_f__Bacteroidales | 20.48 | 18.64 | 16.22 | 16.41 | 21.79 | 5.758 | 0.473 | 0.987 | 0.073 |
| *Prevotella* | 12.79 | 16.90 | 12.48 | 11.08 | 14.76 | 6.015 | 0.635 | 0.878 | 0.919 |
| unclassified_c__Clostridia | 12.06 | 9.77 | 12.03 | 12.56 | 10.14 | 2.121 | 0.063 | 0.534 | 0.664 |
| unclassified_f__Oscillospiraceae | 10.80 | 7.89 | 8.29 | 7.28 | 9.85 | 2.947 | 0.322 | 0.618 | 0.061 |
| unclassified_f__Lachnospiraceae | 5.12 | 4.08 | 5.41 | 4.35 | 5.62 | 1.723 | 0.452 | 0.414 | 0.241 |
| *Methanobrevibacter* | 2.45 | 6.89 | 7.62 | 4.47 | 1.66 | 4.797 | 0.182 | 0.513 | 0.021 |
| *Ruminococcus* | 3.14 | 2.89 | 4.52 | 3.88 | 4.07 | 1.555 | 0.490 | 0.236 | 0.569 |
| unclassified_f__Bacteroidaceae | 2.78 | 3.66 | 2.61 | 3.32 | 3.33 | 1.144 | 0.664 | 0.476 | 0.825 |
| unclassified_c__Bacilli | 3.34 | 2.94 | 1.85 | 4.04 | 2.80 | 1.606 | 0.393 | 0.701 | 0.487 |
| unclassified_o__Eubacteriales | 2.32 | 1.81 | 2.07 | 1.98 | 1.82 | 0.457 | 0.332 | 0.158 | 0.689 |
| unclassified_f__Paludibacteraceae | 1.75<sup>b</sup> | 1.55<sup>b</sup> | 1.42<sup>b</sup> | 1.83<sup>b</sup> | 3.11<sup>a</sup> | 0.905 | 0.004 | 0.003 | 0.004 |
| unclassified_f__Muribaculaceae | 1.70 | 1.52 | 1.32 | 1.86 | 2.26 | 0.990 | 0.616 | 0.374 | 0.174 |
| unclassified_d__Bacteria | 1.06<sup>b</sup> | 1.26<sup>b</sup> | 1.41<sup>b</sup> | 2.10<sup>a</sup> | 1.11<sup>b</sup> | 0.607 | 0.006 | 0.204 | 0.025 |
| unclassified_f__Siphoviridae | 0.71 | 0.75 | 0.85 | 2.31 | 0.68 | 1.292 | 0.117 | 0.394 | 0.264 |
| unclassified_p__Firmicutes | 1.36 | 0.96 | 0.90 | 1.10 | 0.97 | 0.294 | 0.090 | 0.130 | 0.210 |

<sup>1)</sup> SFC 0: Substitution of 0 % ground corn in the basic diet of yak with steam-flaked corn; SFC 25: Substitution of 25 % ground corn in
the basic diet of yak with steam-flaked corn; SFC 50: Substitution of 50 % ground corn in the basic diet of yak with steam-flaked corn; SFC 75: Substitution of 75 % ground corn in the basic diet of yak with steam-flaked corn; SFC 100: Substitution of 100 % ground corn in the basic diet of yak with steam-flaked corn; SEM: standard error of mean.
<sup>2)</sup> Means with different superscript letters in the same row are significantly different ( $p < 0.05$ ).

### 3.6. Relationship between rumen bacteria and fermentation parameters

As shown in Figure 3, the correlation between the relative abundance of rumen bacteria at the genus level and rumen fermentation parameters in yaks is presented. The vertical axis displays the top 20 genera ranked by relative abundance at the genus level, while the horizontal axis represents the rumen fermentation parameters. The propionate content is significantly positively correlated with *Prevotella* and unclassified*_p Bacteroidaceae* (*P* < 0.05); butyrate content is significantly positively correlated with the unclassified*_f Lachnospiraceae* (*P* < 0.05); isobutyrate content is significantly negatively correlated with *Prevotella* and unclassified*_f Acidaminococcaceae* (*P* < 0.05); and the T-VFA content is significantly positively correlated with unclassified*_f Paludibacteraceae* (P < 0.05).

**Fig. 3.**
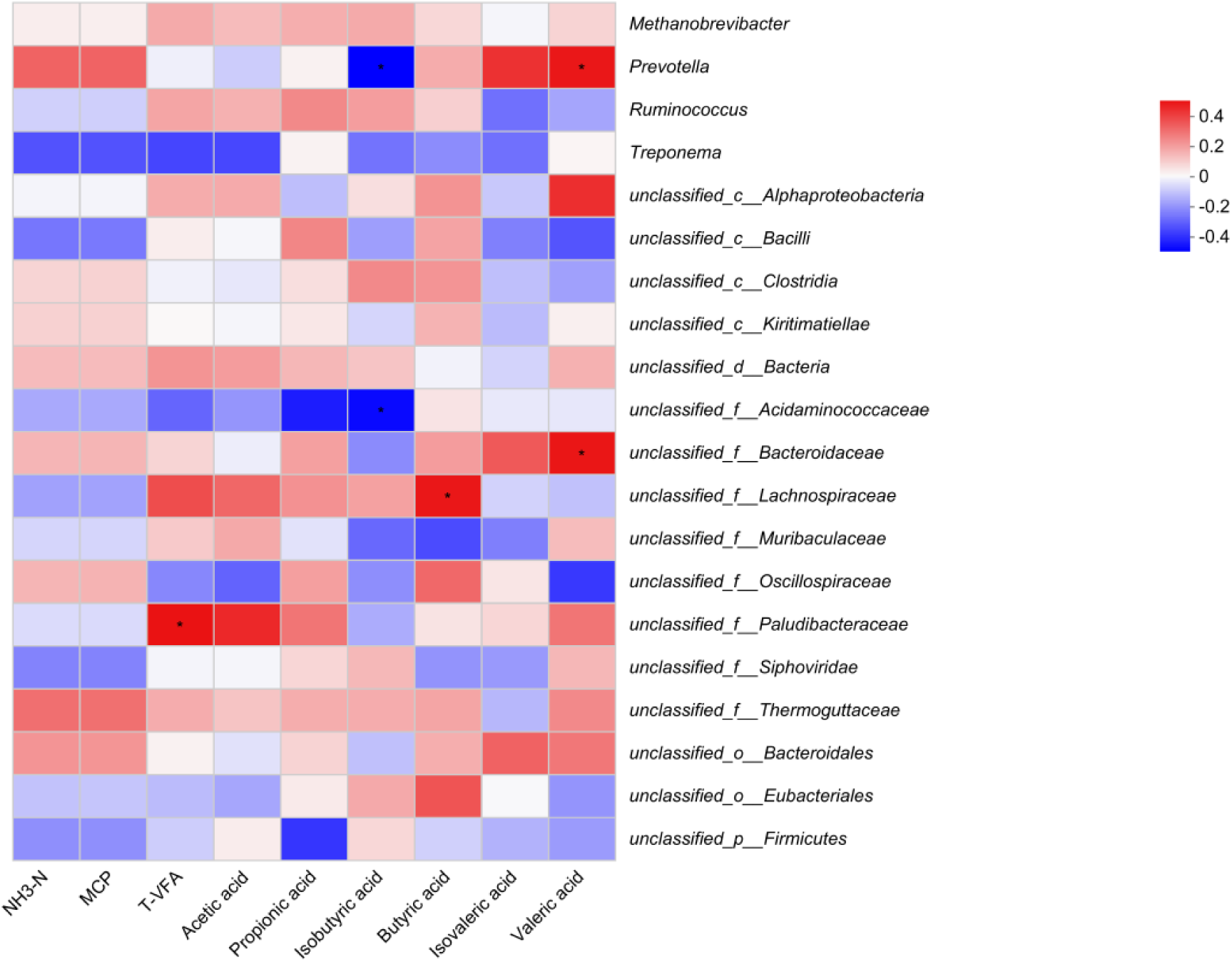
Correlation between relative abundances of bacteria and rumen fermentation parameters. The color and the color intensity of the squares correspond to the direction and strength of the correlation based on the scale to the right. * *p* < 0.05.

### 3.7. CAZy enzyme functional annotation

The abundance of CAZymes was calculated by summing the gene abundances corresponding to each enzyme. A total of 1,611,718 genes were mapped to the CAZy database for this analysis. As shown in Table 7 and Figure 4, the relative abundance of CM in the SFC 0 group was significantly higher than in the other groups (*P* < 0.05), exhibiting a quadratic pattern with a decrease followed by an increase (*P* < 0.05). Carbohydrate esterases (CE) also followed a quadratic trend, initially decreasing and then increasing (*P* = 0.093). The auxiliary activities (AA) in the SFC 50 group showed a trend higher than that in other groups (*P* = 0.051) and the polysaccharide lyases (PL) in the SFC 50 group also exhibited a similar trend (*P* = 0.058).

**Fig. 4.**
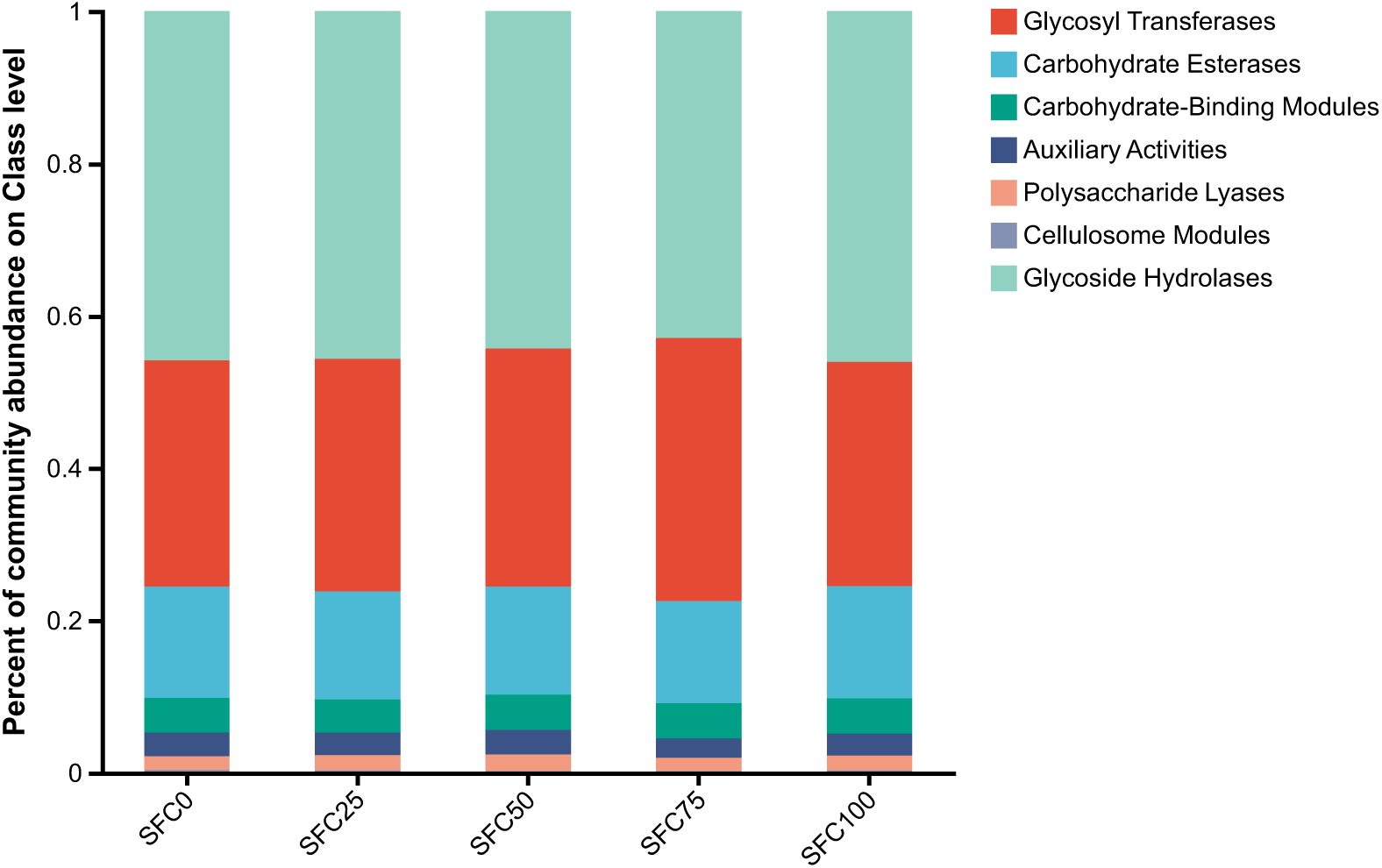
Components of the CAZy enzymes in rumen microbes in different proportions of GC and SFC diet of yaks.

**Table. 7.** Effects of different proportions of GC and SFC diets on the abundance of rumen microbial CAZy enzymes in yaks (%).

| Items | Treatment<sup>1)</sup> |  |  |  |  | SEM | *P* value<sup>2)</sup> |  |  |
| --- | --- | --- | --- | --- | --- | --- | --- | --- | --- |
|  | SFC 0 | SFC 25 | SFC 50 | SFC 75 | SFC 100 |  | Treatment | Linear | Quadratic |
| GH | 45.85 | 44.11 | 44.03 | 42.48 | 45.95 | 0.500 | 0.663 | 0.792 | 0.195 |
| GT | 29.74 | 32.31 | 31.57 | 35.46 | 29.57 | 0.537 | 0.403 | 0.694 | 0.161 |
| CE | 14.60 | 13.77 | 14.13 | 13.16 | 14.68 | 1.319 | 0.257 | 0.768 | 0.093 |
| CBM | 4.57 | 4.44 | 4.66 | 4.59 | 4.65 | 4.403 | 0.974 | 0.696 | 0.901 |
| AA | 3.05 | 3.09 | 3.22 | 2.42 | 2.94 | 5.735 | 0.051 | 0.185 | 0.864 |
| PL | 1.80 | 2.02 | 2.20 | 1.69 | 1.94 | 0.315 | 0.058 | 0.927 | 0.270 |
| CM | 0.38<sup>a</sup> | 0.26<sup>b</sup> | 0.19<sup>b</sup> | 0.19<sup>b</sup> | 0.28<sup>ab</sup> | 0.111 | 0.008 | 0.018 | 0.001 |

<sup>1)</sup> SFC 0: Substitution of 0 % ground corn in the basic diet of yak with steam-flaked corn;
SFC 25: Substitution of 25 % ground corn in the basic diet of yak with steam-flaked corn;
SFC 50: Substitution of 50 % ground corn in the basic diet of yak with steam-flaked corn;
SFC 75: Substitution of 75 % ground corn in the basic diet of yak with steam-flaked corn;
SFC 100: Substitution of 100 % ground corn in the basic diet of yak with steam-flaked
corn; SEM: standard error of mean; GH: glycoside hydrolases; GT: glycosyl transferases;
CE: carbohydrate esterases; CBM: carbohydrate-binding modules; AA: auxiliary activities;
PL: polysaccharide lyases; CM: cellulosome modules.
<sup>2)</sup> Means with different superscript letters in the same row are significantly different ( $p < 0.05$ ).

### 3.8. EggNOG functional protein annotation

As shown in Figure 5, through comparison with the eggNOG database, corresponding NOG annotations were summarized. The most enriched genes were found in translation, ribosomal structure and biogenesis. The second found in the yak rumen were cell wall/membrane/biofilm formation in the eggNOG database (Table 5). As indicated in Table 8, proteins involved in translation, ribosomal structure and biogenesis exhibited a linear decrease with the increase in the substitution ratio of SFC (*P* < 0.05). Proteins with general functional prediction showed a quadratic pattern, initially increasing and then decreasing (*P* < 0.05). Proteins involved in posttranslational modification, protein turnover, chaperones was highest in the SFC 50 group (*P* = 0.041) and showed a quadratic trend of first increasing and then decreasing (*P* < 0.05).

**Fig. 5.**
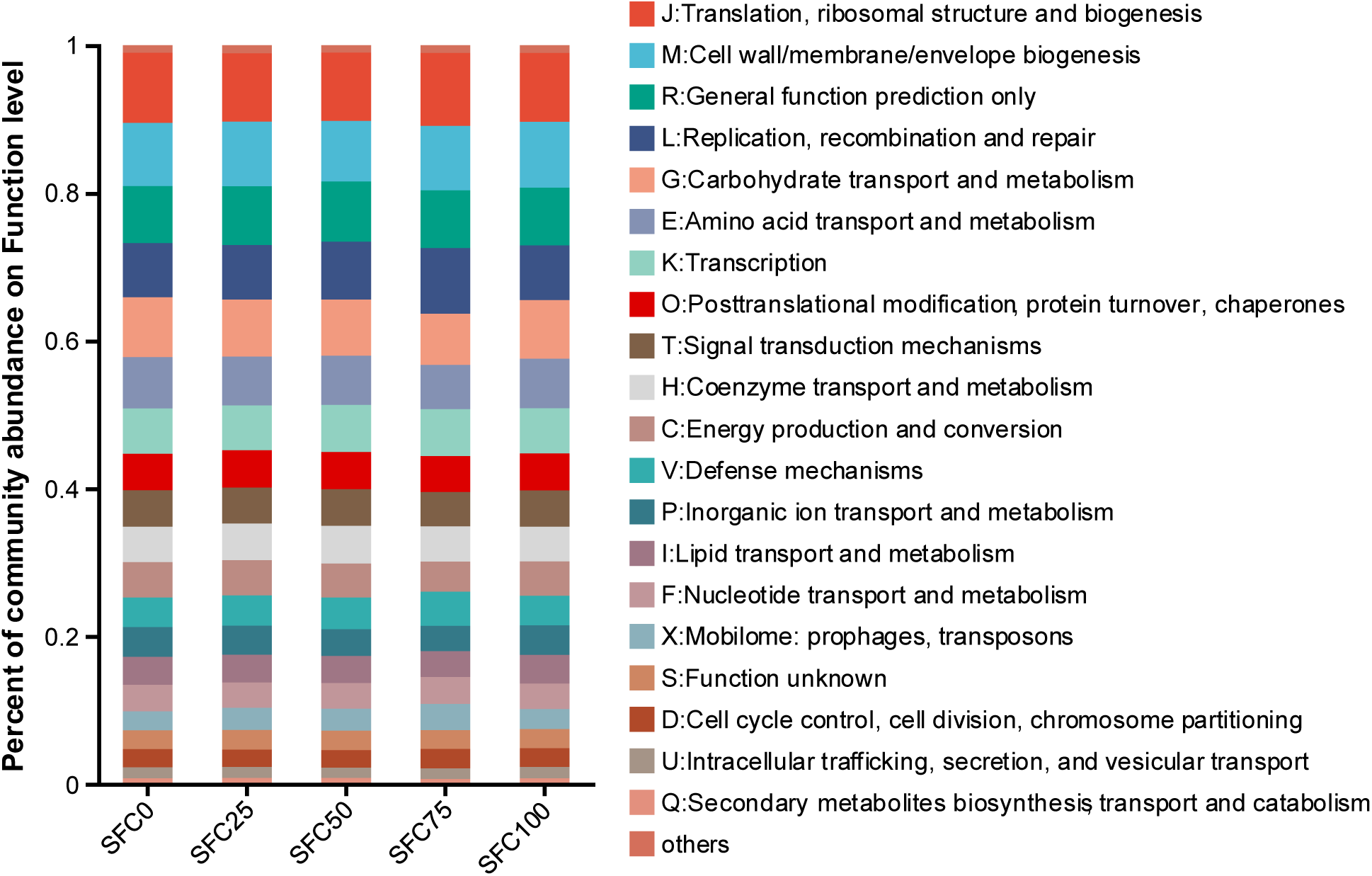
Components of the eggNOG functional protein in rumen microbes in different proportions of GC and SFC diet of yaks.

**Table. 8.**
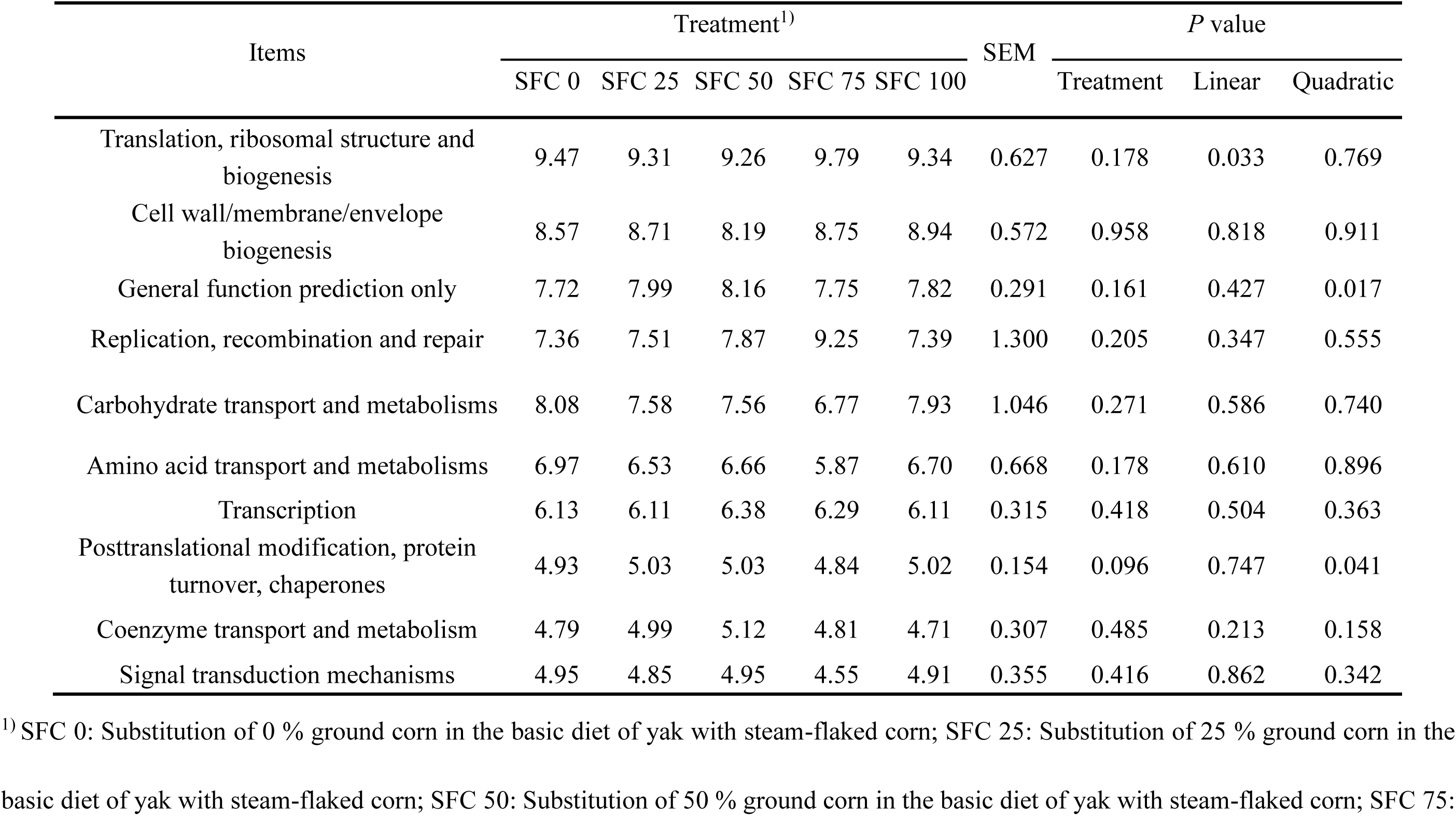
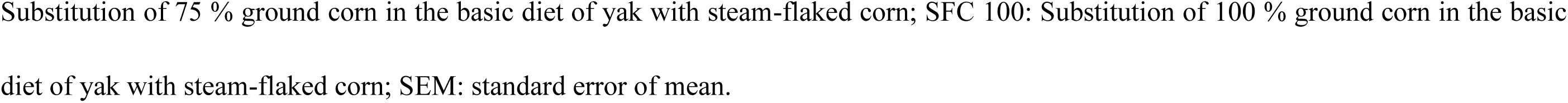
Effects of different proportions of GC and SFC diets on the abundance of rumen microbial eggNOG function protein in yaks (%).

| Items | Treatment<sup>1)</sup> |  |  |  |  | SEM | *P* value |  |  |
| --- | --- | --- | --- | --- | --- | --- | --- | --- | --- |
|  | SFC 0 | SFC 25 | SFC 50 | SFC 75 | SFC 100 |  | Treatment | Linear | Quadratic |
| Translation, ribosomal structure and biogenesis | 9.47 | 9.31 | 9.26 | 9.79 | 9.34 | 0.627 | 0.178 | 0.033 | 0.769 |
| Cell wall/membrane/envelope biogenesis | 8.57 | 8.71 | 8.19 | 8.75 | 8.94 | 0.572 | 0.958 | 0.818 | 0.911 |
| General function prediction only | 7.72 | 7.99 | 8.16 | 7.75 | 7.82 | 0.291 | 0.161 | 0.427 | 0.017 |
| Replication, recombination and repair | 7.36 | 7.51 | 7.87 | 9.25 | 7.39 | 1.300 | 0.205 | 0.347 | 0.555 |
| Carbohydrate transport and metabolisms | 8.08 | 7.58 | 7.56 | 6.77 | 7.93 | 1.046 | 0.271 | 0.586 | 0.740 |
| Amino acid transport and metabolisms | 6.97 | 6.53 | 6.66 | 5.87 | 6.70 | 0.668 | 0.178 | 0.610 | 0.896 |
| Transcription | 6.13 | 6.11 | 6.38 | 6.29 | 6.11 | 0.315 | 0.418 | 0.504 | 0.363 |
| Posttranslational modification, protein turnover, chaperones | 4.93 | 5.03 | 5.03 | 4.84 | 5.02 | 0.154 | 0.096 | 0.747 | 0.041 |
| Coenzyme transport and metabolism | 4.79 | 4.99 | 5.12 | 4.81 | 4.71 | 0.307 | 0.485 | 0.213 | 0.158 |
| Signal transduction mechanisms | 4.95 | 4.85 | 4.95 | 4.55 | 4.91 | 0.355 | 0.416 | 0.862 | 0.342 |

<sup>1)</sup> SFC 0: Substitution of 0 % ground corn in the basic diet of yak with steam-flaked corn; SFC 25: Substitution of 25 % ground corn in the basic diet of yak with steam-flaked corn; SFC 50: Substitution of 50 % ground corn in the basic diet of yak with steam-flaked corn; SFC 75:
Substitution of 75 % ground corn in the basic diet of yak with steam-flaked corn; SFC 100: Substitution of 100 % ground corn in the basic diet of yak with steam-flaked corn; SEM: standard error of mean.

### 3.9. KEGG metabolic pathway annotation

As shown in Figure 6, the relative abundance of KEGG metabolic pathways is ranked in the following order: metabolism, genetic information processing, environmental information processing, cellular processes and organismal systems. As indicated in Table 9, the metabolic pathway of oganismal systems exhibits a quadratic change, initially increasing and then decreasing (*P* < 0.05). The metabolic pathway shows a trend of decreasing first and then increasing (*P* = 0.073). In the SFC 0 group, environmental information processing exhibited a trend that higher than the other treatment groups (*P* = 0.063). Cellular processes showed a quadratic change, initially decreasing and then increasing (*P* = 0.061).

**Fig. 6.**
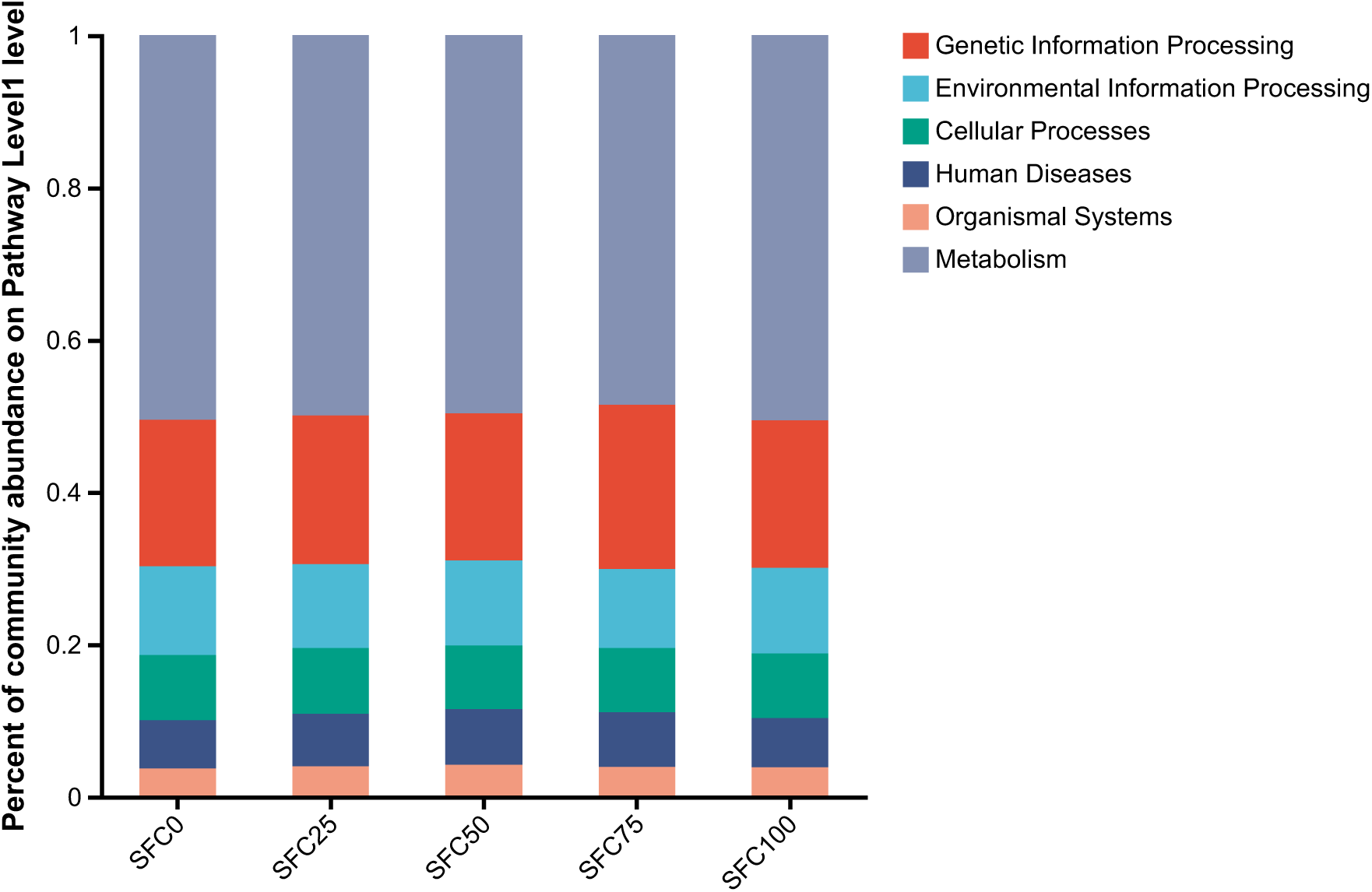
KEGG pathway classification of the rumen microorganisms under different

**Table. 9.** Effects of different proportions of GC and SFC diets on the abundance of rumen microbial KEGG metabolic pathways in yaks (%).

| Items | Treatment<sup>1)</sup> |  |  |  |  | SEM | *P* value |  |  |
| --- | --- | --- | --- | --- | --- | --- | --- | --- | --- |
|  | SFC 0 | SFC 25 | SFC 50 | SFC 75 | SFC 100 |  | Treatment | Linear | Quadratic |
| Metabolism | 50.53 | 49.66 | 49.58 | 48.51 | 50.56 | 1.861 | 0.473 | 0.987 | 0.073 |
| Genetic Information Processing | 19.23 | 19.81 | 19.46 | 21.93 | 19.46 | 1.994 | 0.635 | 0.878 | 0.919 |
| Environmental Information Processing | 11.66 | 10.96 | 11.13 | 10.09 | 11.20 | 0.945 | 0.063 | 0.534 | 0.664 |
| Cellular Processes | 8.55 | 8.64 | 8.36 | 8.39 | 8.49 | 0.372 | 0.322 | 0.618 | 0.061 |
| Organismal Systems | 3.70 | 4.01 | 4.17 | 3.89 | 3.87 | 0.423 | 0.182 | 0.513 | 0.021 |

<sup>1)</sup> SFC 0: Substitution of 0 % ground corn in the basic diet of yak with steam-flaked corn; SFC 25: Substitution of 25 % ground corn in the basic diet of yak with steam-flaked corn; SFC 50: Substitution of 50 % ground corn in the basic diet of yak with steam-flaked corn; SFC 75: Substitution of 75 % ground corn in the basic diet of yak with steam-flaked corn; SFC 100: Substitution of 100 % ground corn in the basic diet of yak with steam-flaked corn; SEM: standard error of mean.

## 4. Discussion

Yaks, as the primary livestock species in the western Sichuan Plateau, face unique challenges in farming due to the region’s distinctive geographical and climatic conditions, making yak farming development more difficult than for other livestock species (Shah A M et al.,2023). Issues such as the feeding cycle and the grass-livestock conflict have made optimizing the use of major feed ingredients in yak diets a key breakthrough in improving farming efficiency. The feed ingredients in the diet of yak play a critical role in its growth and development, and enhancing the nutritional level of the diet can significantly promote yak growth (Xu T. et al., 2017). Antonio et al. replaced GC with SFC in the diet of Nellore bulls (Arriola Apelo et al., 2014), finding that the SFC group had a 7% reduction in feed intake, but improvements in ADG and feed conversion efficiency. In this experiment, the ADG of yaks in all groups showed a quadratic trend of first increasing and then decreasing. This suggests that replacing 50% of GC with SFC in yak diets results in optimal growth performance, effectively improving ADG and FCR.

Various biochemical indicators in serum can be used to assess the health status of yaks as well as the digestion and metabolism of nutrients in their diets. TG is one of the key indicators of energy and fat metabolism in the body. In this experiment, the serum TG levels in the SFC 100 group were significantly higher than those in other treatment groups, and exhibited a quadratic trend of first decreasing and then increasing as the proportion of SFC replaced ground corn. This suggests that replacing ground corn with SFC in the diet affects yak fat metabolism, which is consistent with the findings of Wu Shufeng (Wu S.F. et al., 2021). GLU reflects energy metabolism in the body (Reeds, P.J. et al., 2000) GLU is the primary direct energy source in animal metabolism, and changes in serum GLU concentration reflect the dynamic balance of the breakdown, absorption, and transport of carbohydrates such as starch and cellulose in the diet, as well as the balance between energy levels, glucose synthesis, and consumption. In this experiment, the serum GLU concentration in the SFC 50 group tended to be higher than that in other treatment groups, suggesting a more balanced energy metabolism in the SFC 50 group, which may promote the synthesis of glycogen and increase serum GLU levels. PYR is a key intermediate product in the metabolism of carbohydrates, fats, and proteins. Under anaerobic conditions, it is converted into lactic acid, whereas in the presence of sufficient oxygen, it enters the mitochondria to be oxidized into carbon dioxide and water (Reeds, P.J. et al., 2000). In this experiment, the serum PYR levels in the SFC 50 group tended to be higher than those in other treatment groups, indicating that anaerobic metabolism was reduced to some extent, improving energy utilization, which is consistent with previous research findings (Finazzi and Arosio, 2014).

Rumen fermentation parameters are important indicators for assessing the stability of the rumen environment and the extent of rumen fermentation. In this experiment, no significant changes in NH_3_-N and MCP were observed among the treatment groups, which is consistent with the findings of Malekkhahi et al. (Malekkhahi, et al. 2021;Spears et al. ^2^00^4^). The levels of NH_3_-N and MCP in all groups were within the appropriate concentration range; however, the SFC 50 group exhibited higher values of NH3-N and MCP compared to the other groups. This suggests that under these conditions, rumen microbial fermentation may be more favorable, leading to more efficient utilization of energy by the rumen microbes. Rumen microbes ferment the carbohydrates in the diet to produce a large quantity of VFA, which not only provide 70-80% of the animal’s energy but also serve as carbon skeletons for amino acids (Li et al., 2021). As the most important products of carbohydrate degradation by rumen microbes, T-VFA content is often directly related to the animal’s production performance (Njokweni et al., 2021). Acetic acid, propionic acid, and butyric acid are the three main VFAs, accounting for more than 95% of the total, with acetic acid being the main precursor for milk fat synthesis and propionic acid being involved in gluconeogenesis to produce glucose (Domenico et al., 2007). VFAs affect the pH of the rumen, which in turn determines the ability of rumen microbes to digest and absorb nutrients. Optimizing the ratio of acetic acid to propionic acid and promoting a fermentation pattern dominated by propionic acid can provide more usable energy to the animal, thereby optimizing rumen fermentation (Hansen et al., 2009). In the in vitro gas production experiment by Zhang Ting et al. (Spears, et al., 2004), the SFC group showed significantly higher T-VFA and propionic acid content compared to the regular corn group, which notably improved rumen fermentation characteristics. Liu Ping et al. found in their study on Dehong hybrid buffalo that SFC significantly increased propionic acid levels and reduced the ratio of acetic acid to propionic acid (Gou et al., 2018). In this experiment, as the proportion of SFC replaced ground corn, the propionic acid content in the yak rumen fluid significantly increased, while acetic acid and total VFA content increased linearly, which is consistent with previous research results. The SFC 50 group had the lowest ratio of acetic acid to propionic acid, suggesting that this condition resulted in a more favorable rumen fermentation pattern, providing more usable energy to the yaks, which is in line with the growth performance results mentioned earlier.

Rumen microbes, including bacteria, protozoa, fungi and other microorganisms, play a crucial role in breaking down proteins, starches, and celluloses, thus providing an important energy source for the host (Park et al. 2020). Therefore, a comprehensive understanding of the rumen microbial community is essential for the digestion and utilization of nutrients in ruminant animals. Research has shown that the source and level of dietary energy can alter the rumen microbial community in lactating cows (Takizawa et al., 2020). Specific functional enzymes, such as proteases, amylases, and pectinases, produced by rumen microbes can break down nutrients in the animal’s diet into small peptides and sugars (Kamra, 2005;Shi et al., 2014). A large body of research indicates that changes in the rumen microbial community can affect energy efficiency (Gan et al., 2023). Previous studies have demonstrated that the dominant bacterial phyla in the rumen are *Firmicutes* and *Bacteroidetes* (Hu et al., 2020;Xu et al., 2003). Similarly, in this study, the primary dominant phyla in the yak rumen across all groups were *Firmicutes*, *Bacteroidetes*, *Euryarchaeota*, and *Proteobacteria*. *Firmicutes* promote energy absorption by the host through the production of glycoside hydrolases. Based on 16S rDNA gene sequencing, *Bacteroidetes* can metabolize substances like steroids and bile acids, promoting the absorption of polysaccharides (Gan et al., 2023). The role of *Uroviricota* in the digestive tract remains poorly understood; however, previous studies have shown that *Uroviricota* and *Actinobacteria* typically exhibit opposite trends in abundance (Dominika et al., 2024), which was also observed in this experiment. The relative abundance of *Uroviricota* showed a quadratic increase followed by a decrease, whereas the relative abundance of *Actinobacteria* exhibited a quadratic decrease followed by an increase. The fungal phylum *Planctomycetota* encodes numerous Cazymes, demonstrating significant hydrolytic potential for the degradation of complex polysaccharides (Opdahl et al., 2018). In the SFC 50 group, the relative abundance of *Planctomycetota* was significantly higher than that in other groups, offering new insights into its potential for the degradation of total carbohydrates.

At the genus level, in this study, the *Bacteroides*, *Prevotella*, *Clostridium*, *unclassified_f Oscillospiraceae*, and unclassified*_f Lachnospiraceae* dominated the microbial community, consistent with findings from previous research on various ruminant species (Dai et al., 2021;Hua et al., 2022;Huang et al., 2012;Wei et al., 2021). *Methanobrevibacter* is a major methanogen in the rumen of ruminants. In this study, the relative abundance of *Methanobrevibacter* exhibited a quadratic increase followed by a decrease, which aligns with previous in vitro studies showing an increase in *Methanobrevibacter* relative abundance and higher gas production in the SFC group (Malik et al., 2021)., The *Bacteroidales* plays a role in the degradation of proteins and amino acids, and some *Bacteroidales* are capable of non-glycolytic metabolism (Li et al., 2021). The relative abundance of *Bacteroidales* in this study showed a quadratic decrease followed by an increase, which is consistent with the results of a previous study that used steam-treated grains to feed ponies (Ley et al., 2006). Some studies have suggested that unclassified*_d Bacteria* are associated with NO emissions (Ley, et al., 2006), and they represent the dominant microbial community in environments with reduced nitrogen and phosphorus levels (Li et al., 2023). In the present experiment, the relative abundance of unclassified*_d Bacteria* was highest in the SFC 75 group. *Paludibacteraceae* are capable of transporting glucose, xylose, and maltose (Wang et al., 2023), and in this experiment, the relative abundance of unclassified*_f Paludibacteraceae* in the SFC 100 group was significantly higher than that in the other groups, showing a quadratic increase followed by a decrease. This suggests that replacing GC with SFC in the yak diet can enhance the ability to degrade carbohydrates of rumen microorganisms. Additionally, the relative abundance of unclassified*_c Clostridia* significantly decreased in this experiment, which is consistent with the findings of Wang et al. in goats (Rice et al., 2012).The relationship between The relative between the abundance of rumen bacteria at the genus level and rumen fermentation parameters has been extensively studied. Previous research has found a strong correlation between the content of valerate and the relative abundance of *Prevotella* (Hinnebusch et al., 2002), which is consistent with the findings of the present study. In this study, valerate content was also positively correlated with unclassified*_f Bacteroidaceae*, a result similar to that observed in previous research on microbial communities in the colon of swine (Van den Abbeele et al., 2022). Furthermore, unclassified*_f Paludibacteraceae* efficiently utilize energy and amino acids to produce higher amounts of VFA, and their abundance showed a significant positive correlation with the T-VFA.

The CAZy enzymes produced by the rumen microbiome during the breakdown of carbohydrates are involved in various biological processes in the body, including carbohydrate metabolism, protein glycosylation, and degradation processes (Gong et al., 2020). Research by Gong et al. (Jiang et al., 2022) indicated that the fecal microbiome of yaks varies according to the types of food they consume. Due to seasonal variations in feed composition, the relative abundance of CAZy enzymes in the yak rumen also exhibits seasonal changes (Drula et al., 2021). In this study, the relative abundances of AA and PL in the SFC 50 group were higher than in other treatment groups, while CM was significantly lower and showed a quadratic curve, initially decreasing and then increasing. PL is involved in the breakdown of polysaccharides containing uronic acids, generating unsaturated hexadecenoic acid and a new reducing end (Mhatre, N. et al., 2025). In this trial, the relative abundances of PL and AA were highest in the SFC 50 group, which may promote rumen bacterial degradation of carbohydrates and enhance the activity of vitality-supporting enzymes. CM is a self-assembled multi-enzyme complex capable of efficiently degrading plant cell walls, particularly in the breakdown of lignocellulose (Lee, et al., 2023). In this study, the relative abundance of CM was highest in the SFC 0 group, which may be related to the higher fiber content in the crushed corn.

EggNOG is a bioinformatics database and tool that provides evolutionary information about genes and proteins. In the EggNOG database, functional proteins refer to groups of proteins with similar structures and functions, identified through the comparison and analysis of proteins from different species. This similarity is typically determined by aligning protein sequences and structures. By performing these alignments and classifications, a better understanding of the function of proteins in organisms and their evolutionary changes can be achieved (Ramazi et al., 2021). In the present study, the EggNOG functional proteins in the yak rumen microbiome involved in translation, ribosomal structure and biogenesis showing a linear variation, indicating that as the content of flaked corn increased, protein generation also changed. The functional proteins involved in posttranslational modification, protein turnover, chaperones in the SFC 50 group showed a trend of higher than that in other treatment groups, with a quadratic pattern of first increasing and then decreasing. These proteins can alter the chemical properties of proteins, such as phosphorylation, methylation, acetylation, thereby regulating the structure and function of proteins. Protein turnover controls the rate of protein synthesis and degradation, maintaining a dynamic balance of protein levels within cells (Marchingo, J.M. et al., 2022). The KEGG Pathway database contains extensive biological pathway information. Each pathway consists of a series of interacting molecules and reactions that describe the interactions and regulatory relationships between these molecules (Yang, L. et al., 2022). In this study, we primarily analyzed the category level. Among these, metabolism pathways are linked by a series of interacting compounds and enzymes. These pathways describe the processes and regulation of different metabolic processes in organisms. Each pathway involves a specific enzyme that catalyzes the conversion of substrates into products. The classification of metabolism pathways is typically based on different aspects of metabolism in organisms, such as energy metabolism, carbohydrate metabolism, amino acid metabolism, and others (Zhang, K. et al.,2023). In this study, metabolism pathways showed the highest enrichment. Environmental information processing pathways refer to a series of biological pathways that involve the perception, transmission, and response to external environmental signals within an organism. These pathways enable organisms to adapt to environmental changes, maintain homeostasis, and make appropriate physiological and behavioral responses to environmental factors (Chen L et al., 2017). In the SFC 0 group, the environmental information processing pathways were significantly higher than in other treatment groups. Cellular process pathways involve the regulation, execution, and modulation of various biological processes within the cell, including cell cycle regulation, apoptosis, cell signaling, and cell differentiation. In this study, cellular process pathways exhibited a quadratic trend of first decreasing and then increasing.

## 5. Conclusions

This study found that replacing 50% of the GC in the diet with SFC increased the ADG of the yaks, as well as the serum GLU and PYR levels. Furthermore, substituting SFC for GC in the diet no matter improved the shift of rumen fermentation towards propionate-type fermentation but also optimized the rumen microbial community structure. Meanwhile, it influenced the rumen CAZy enzymes and eggNOG functional proteins and enhanced the efficiency of carbohydrate and protein utilization in yaks. In conclusion, the results above suggest that replacing 50% of the GC with SFC in the diet is an effective feeding strategy for fattening yaks.

## Acknowledgements

This work was supported by the National Modern Agricultural Industrial Technology System Sichuan Beef Cattle Innovation Team Building Project (sccxtd-2023-13); the research on Key Technology and Integrated Application of efficient yak breeding (20230044).

## Declaration of competing interest

The authors declare that they have no conflict of interest.

## Notes

Financial support: This study supported by the National Modern Agricultural Industrial Technology System Sichuan Beef Cattle Innovation Team Building Project (sccxtd-2023-13); the research on Key Technology and Integrated Application of efficient yak breeding (20230044).

### Competing Interest Statement

The authors have declared no competing interest.

